# Frequent spontaneous structural rearrangements promote rapid genome diversification in a *Brassica napus* F1 generation

**DOI:** 10.1101/2022.06.27.497715

**Authors:** Mauricio Orantes-Bonilla, Manar Makhoul, HueyTyng Lee, Harmeet Singh Chawla, Paul Vollrath, Anna Langstroff, Fritz J. Sedlazeck, Jun Zou, Rod J. Snowdon

## Abstract

In a cross between two homozygous *Brassica napus* plants of synthetic and natural origin, we demonstrate that novel structural genome variants from the synthetic parent cause immediate genome diversification among F1 offspring. Long read sequencing in twelve F1 sister plants revealed five large-scale structural rearrangements where both parents carried different homozygous alleles but the heterozygous F1 genomes were not identical heterozygotes as expected. Such spontaneous rearrangements were part of homoeologous exchanges or segmental deletions and were identified in different, individual F1 plants. The variants caused deletions, gene copy-number variations, diverging methylation patterns and other structural changes in large numbers of genes and may have been causal for unexpected phenotypic variation between individual F1 sister plants, for example strong divergence of plant height and leaf area. This example supports the hypothesis that spontaneous *de novo* structural rearrangements after *de novo* polyploidization can rapidly overcome intense allopolyploidization bottlenecks to re-expand crops genetic diversity for ecogeographical expansion and human selection. The findings imply that natural genome restructuring in allopolyploid plants from interspecific hybridization, a common approach in plant breeding, can have a considerably more drastic impact on genetic diversity in agricultural ecosystems than extremely precise, biotechnological genome modifications.

## Introduction

Genetic and genomic diversity plays a key role in plant adaption to environmental changes. Plant breeding exploits such diversity to develop new varieties adapted to new growth environments or biotic and abiotic challenges (Louwaars, 2018). In many crops the exploitation of heterosis through hybrid breeding, achieved via expansion and differentiation of genetic and genomic diversity into distinct heterotic pools, has been crucial to breeding success (Labroo et al., 2021). However, both in hybrid and open-pollinated crops, genetic bottlenecks due to inbreeding or targeted trait selection are a constant obstacle in breeding programs (Hickey et al., 2019). Furthermore, numerous important crop species arose through allopolyploidization, a process that is sometimes described as an evolutionary dead-end because of the severe genetic bottleneck posed by small numbers of founders involved to allopolyploidization events (Mayrose et al. 2015, Soltis, et al. 2014). Detailed analysis of genetic and genomic diversity in crop germplasm pools and breeding materials has become an important technique to identify and exploit diversity for breeding, and in recent decades genome-wide assays of sequence polymorphisms have become a relevant tool for genomic selection and gene discovery in crops. High-throughput sequencing technologies also provide a means to investigate the consequences and impact of polyploidization on genome structure and genome-scale diversity (Samans et al., 2017).

High-throughput genetic analysis of large plant populations normally implements single nucleotide polymorphism (SNP) arrays or genotyping-by-sequencing techniques which deliver low-cost, relatively simple and meanwhile reasonably standardized datasets for extrapolation or imputation of genome-wide DNA sequence patterns. New SNP variants can also arise spontaneously during mitosis and meiosis. Mutations of this kind occur in plants at different rates depending on the genome size and ploidy. For example, in *Arabidopsis thaliana*, between 1 and 5 *de novo* intra-varietal mutations have been shown to occur per generation, whereas higher rates are found in rice and other plants with larger genomes (Singer et al., 2021). However, such estimates do not account for internal and external variables that can induce mutation, like cell age, epigenetics or temperature (Schoen and Schultz, 2019).

Meiosis also generates other forms of genomic diversity, not only by recombination through crossovers (CO), but also via meiotic mutations in gametes due to errors in the repair of DNA double-strand breaks (DSB). In particular, non-homologous end-joining (NHEJ) (Kuo et al., 2021; Wang et al., 2021) can lead to a variety of *de novo* genomic rearrangements (Cai and Xu, 2007). According to the theoretical framework of Mendel’s laws of inheritance (Mendel, 1866), meiotic mutations should be inherited in heterozygous form by all F1 offspring whose parents are highly homozygous. In plants, examples of non-Mendelian inheritance have been identified, for example template-directed extra-genomic sequence insertions in *A. thaliana* (Lolle et al., 2005) or selfish genetic elements leading to segregation distortion in rice (Yu et al., 2018).

In this study, we used long read sequencing to detect unexpected inheritance patterns across a set of F1 sister plants developed from a cross between the genetically diverse *B. napus* accessions Express 617 and *B. napus* G3D001. Express 617 is an inbred (F11), natural winter-type oilseed rape that has been widely used in genetic and genomic analyses (Lee et al., 2020), while G3D001 is an advanced homozygous synthetic *B. napus* line derived from crosses between *B. napus, B. rapa* (AA, 2n = 20) and *B. carinata* (BBCC, 2n = 34), as described by Zou et al. (2018).

The important oilseed crop plant *Brassica napus* (genome AACC, 2n = 38) originated only very recently (Chalhoub et al., 2014) from interspecific crosses between the closely related diploid progenitors *B. rapa* and *B. oleracea* (CC, 2n = 18). Its recent origin from a limited number of founders and intensive selection for important seed quality characters in the past several decades represent extreme genetic bottlenecks. Paradoxically, despite its narrow genetic basis, *B. napus* has very quickly become one of the world’s most important oilseed crops and profited from tremendous breeding success. Its importance as an oilseed crop and its closeness to *A. thaliana* have made it an interesting model for polyploid crop evolution. A striking feature in this context is the broad prevalence of genomic structural variations (SV), first discovered as large-scale homoeologous chromosome exchanges in genetic mapping studies (Song et al., 1995) and later confirmed by fluorescence *in situ* hybridization (Xiong et al., 2011), transcriptome-based visualization (He et al., 2017), or whole-genome assembly and genome resequencing (Chalhoub et al., 2014; Samans et al., 2017; Hurgobin et al., 2018). In *B. napus*, homoeologous non-reciprocal translocations (HNRT) and other homoeologous recombinations between highly similar chromosomes are particularly prevalent and often very large in synthetic *B. napus*, but also commonly found in naturally-derived accessions (Chalhoub et al., 2014; Higgins et al., 2018).

Samans et al. (2017) postulated that elevated frequencies of HNRT in early generations after *de novo* polyploidization could be an important driver for novel genetic variation to overcome the allopolyploidy bottleneck in evolution and breeding. Subsequently, Higgins et al. (2018) used allele presence-absence data from the *Brassica* 60K SNP array (Mason et al., 2017) to detect *de novo* homoeologous recombination events in test-cross families derived from a panel of 11 *B. napus* cultivars, demonstrating for the first time that *de novo* HNRT indeed generates novel, unexpected genetic diversity during *B. napus* breeding. Using transcriptome sequencing, Lloyd et al. (2018) observed *de novo* homoeologous exchanges between individual plants of the same inbred *B. napus* cultivar.

In recent years, advances in long read sequencing techniques achieved highly accurate resolution of homoeologous chromosome regions in allopolyploid genome assemblies (Lee et al., 2020; Rousseau-Gueutin et al., 2020). Furthermore, resequencing using long-read sequencing provided a technical platform for accurate, routine detection of SV in complex plant genomes, including that of *B. napus* (Mahmoud et al., 2019; Yuan et al., 2021). Unexpectedly, surveys of genome-wide SV extent and patterns using Oxford Nanopore Technology (ONT) and/or Pacific Biosciences (PacBio) long read sequencing techniques suggested that all individual *B. napus* accessions carry many small-to medium-scale SV events within genic regions, with direct functional implications (Gabur et al., 2020; Vollrath et al., 2021a) and a potentially major role as drivers of genetic diversity and phenotypic adaptation (Chawla et al., 2021).

Here, we used ONT long reads to sequence 12 F1 sister plants of *B. napus* derived from a single cross between two strongly homozygous parents. As in the F1 test cross families investigated previously by Higgins et al. (2018), the 12 sister hybrids in our study would be expected to be genetically uniform according to Mendel’s law of uniformity were investigated in detail on a genome-wide scale for high-resolution detection of spontaneous genomic rearrangements that were not observed in the genomes of the two parents. This was achieved by combining putative SV alleles with read coverage information after alignment and SV calling. Homoeologous exchanges were linked to large structural rearrangements and methylation patterns were predicted from ONT reads to assess the putative transcriptomic and epigenomic effects from observed spontaneous mutations. We confirmed that genome-wide rearrangements derived from a recent allopolyploid plant can give rise to vastly different new genetic variants in just a single generation. These observations add to the growing body of evidence that homoeologous exchanges can lead to rapid and ongoing diversification of allopolyploids crops during evolution and breeding, despite the enormous bottleneck of a spontaneous interspecific hybridization.

## Materials and Methods

### Development of F1 sister plants

Parental lines G3D001 (Zou et al., 2018) and Express 617 (Lee et al., 2020) were sown in Hawita propagation substrate “F.-E. Typ P” (Hawita Gruppe GmbH) and placed in a greenhouse chamber in Giessen, Germany with a 16:8 hour light/dark photoperiod and an average temperature and relative humidity of 5 °C and 65% RH, respectively. After 6 weeks, Express 617 seedlings were transferred to a separate chamber for 10 weeks of vernalization at 5°C and 65% RH and 16:8 light/dark photoperiod. The seedlings were then re-transferred inside the greenhouse and grown along with G3D001 under the same conditions. Pollen from a single plant of the paternal parent G3D001 was used to pollinate a single plant of the maternal parent Express 617 after emasculating immature maternal flower buds with alcohol sterilized tweezers. The crosses were immediately labelled and covered with a plastic bag to prevent cross-pollination.

### Material sampling and phenotyping

F1 seeds were harvested, sown and the resulting F1 sister plants were grown and vernalized under the same conditions as described for the cross parents. Twelve sister F1 plants, along with one plant each from the maternal and paternal parents, were transplanted to 120 litre large plant containers (Hohmann et al., 2016) containing a homogenized 60/40 sand-soil mix and grown side-by-side, alongside one plant per parental line, under semi-controlled conditions in a tunnel greenhouse at Rauischholzhausen, Germany to ensure a uniform growing environment in a large soil volume for all 14 plants. All plants in the greenhouse unit were phenotyped with a 3D PlantEye Dual-Scanner F500 (Phenospex) for 11 weeks from the seedling stage to full flowering stage to evaluate their morphological uniformity, and an identical watering regime was applied to all plants. The second or third youngest leaf of each plant was harvested at 11:00 am on the same day and then frozen with liquid nitrogen and stored at -80°C until further processing. Leaves from each F1 plant and the two parental plants used as crossing parents were subsequently ground in liquid nitrogen using a sterilized mortar and pestle.

### DNA isolation and long read sequencing

High-molecular-weight (HMW) DNA was extracted following a previously described protocol (Chawla et al., 2021). DNA quality and length were evaluated with a Nanodrop spectrophotometer (Thermo Fisher), a Qubit 2.0 fluorometer (Thermo Fisher) and gel electrophoresis. Libraries were prepared using SQK-LSK109 ligation sequencing kits (Oxford Nanopore Technology) and were afterwards loaded on Oxford Nanopore R9.4.1 flow cells in a MinION sequencing device (Oxford Nanopore Technology) for the G3D001 plant used to develop the F1 sister plants, and in a PromethION (Oxford Nanopore Technology) sequencing platform for all other samples.

### Base calling and long read data filtering

Raw electrical signals from plants grown in Rauischholzhausen were base-called using Guppy Basecaller v.4.0.11 (Oxford Nanopore Technology), in a virtual machine operating with Ubuntu 20.04.1 LTS with two NVIDIA Tesla 4 TU104GL (NVIDIA Corporation) Graphic Processor Units (GPU) and using the following options: *--device cuda: 0,1:50% --kit SQK-LSK109 -- num_callers 16 -- disable_pings* and *--flowcell FLO-PRO002*. Reads from the G3D001 plant used as pollen donor for the F1 generation were base-called with the same settings except for *FLO-MIN106* with Guppy Basecaller v.5.0.7. Only reads with a quality score above 7 and length above 5000 nucleotides were kept using NanoFilt v.2.8.0 (de Coster et al., 2018). The filtered library quality was evaluated with NanoStat v.1.5.0 (de Coster et al., 2018) and genome-wide coverages were estimated.

### Structural variation calling

Filtered long reads from G3D001 and all F1 sister plants were aligned against the Express 617 reference genome (Lee et al., 2020) using minimap2 v. 2.20 (Li, 2018) *map-ont* function with *-ax* settings. The output file of each alignment was then filtered using samtools v.1.12 (Li et al., 2009) *view* function, so that only reads with an alignment score above 50 were kept. Mid-sized structural variations longer than 30 bp and supported by at least 25 reads were called using sniffles v.1.0.12 (Sedlazeck et al., 2018). Only insertions and deletions detected through aligned and/or split reads, having precise breakpoints and with resolved lengths for insertions were kept and merged with the forced calling pipeline from SURVIVOR v.1.0.7 (Jeffares et al., 2017) to allow SV comparison across samples. Moreover, only SVs where G3D001 had a homozygous alternate allele, smaller than 50 kbp and without miscalled alleles from any sample were selected. Furthermore, SVs which had less than 90% of reads supporting the predicted allele were discarded, in order to reduce false positives due to residual heterozygosity. SVs having different alleles across F1 sister plants were identified based on the variant calling files and further visualized with the Integrative Genomics Viewer (IGV) tool (Robinson et al., 2011). Selected insertions were assessed with polymerase chain reaction (PCR) and agarose gel electrophoresis using DNA from all 12 F1 sister plants and their two parents.

### Detections of large genomic rearrangements above 1 Mbp

1Mbp windows were prepared for each chromosome using bedtools (Quinlan and Hall, 2010) *makewindows* and *coverage* functions. 1Mb window coverages were then combined with the allele information and position of SVs that were putatively different across F1 sister plants in tab-delimited files based on the allele type: homozygous reference, homozygous alternate and heterozygous. The SV coverage and allele genotype were visualized using the circlize package (Gu et al., 2014). Regions larger than 1 Mbp in which one or more F1 genotype showed no heterozygous SV alleles were further visualized with IGV to estimate the rearrangement start and end based on the genomic positions at which the coverage halved. Due to the large memory requirements to display coverages from all genotypes in large regions at the same time, plots for each genotype were saved as images separately instead and then merged with GIMP for easier display. Moreover, the number of genes within each large rearrangement were found by using the *intersect* function from bedtools against the Express 617 gene annotation. The genes functions were assessed by blasting their complementary DNA (cDNA) sequences with BLASTn (Altschul et al., 1990) against the Araport 11 *A. thaliana* representative gene model cDNA sequences (Cheng et al., 2017). Only the hit with lowest e-value was kept, on the condition that it had an e-value lower than 1 × 10^−4^, no opening gaps and a percentage of identity equal or higher than 90%.

### Centromere prediction

Centromeres were predicted to define their distance to detected large genomic rearrangements. Briefly, two centromere-specific repeat sequences CentBr1 (GenBank accession CW978699) and CentBr2 (GenBank accession CW978837) were used to estimate the approximate positions of the centromeric regions for each chromosome. The methods were based on (Mason et al., 2016), where the two sequences where aligned to the Express 617 chromosomal assembly using BLASTn (Altschul et al., 1990) with a cut off of at least 90% sequence similarity. Since these sequences are satellite repeats that are not limited to only centromeric regions, further refinement of the centromeric positions was required. Using the approximate position range obtained through the alignment results representing the centromere boundaries, we traced them back to the scaffolding process of Express 617 assembly to find breakpoints. Breakpoints were defined as the positions where two non-overlapping scaffolds were merged together through genetic maps (Lee et al., 2020). These breakpoints were then set as the refined version of the centromeric boundaries and were used to estimate the relative position of large rearrangements to centromeres.

### Identification of homoeologous exchanges

Recent assemblies from *B. oleracea* (Lv et al., 2020) and *B. rapa* (Zhang et al., 2018) were concatenated and used for homoeologous exchange identification. Homologous gene pairs between the A and C subgenomes were located with inparanoid v.4.2 (O’Brien et al., 2005) using bootstrap, a BLOSUM80 (BLOcks SUbstitution Matrix) and an initial cut-off score of 60. Inparalogs with a similarity score equal or greater than 70 were selected for each gene. Only pairs with the highest similarity score were kept and only the first reported homologous gene pair was selected in cases where two or more gene pairs had the same similarity score. Quality filtered long reads from plants grown in Rauischholzhausen were aligned with minimap2 against the concatenated *B. napus* reference. Coverage across chromosomes was calculated using the *bamtobed* and *genomecov* functions from bedtools and used as input in a modified deletion-duplication pipeline previously described (Stein et al., 2017). Briefly, outlier regions with a coverage above 150 were discarded and segments equal or larger than 25000 bp, with a coverage that deviated by at least one standard deviation above or below the mean coverage, were called as duplication or deletion, respectively. Those segments in which a gene homolog was deleted and its reciprocal homolog was duplicated in a homoeologous chromosome, were considered as putative non-reciprocal homoeologous exchanges; these were further searched within large genomic rearrangements (> 1 Mbp) to determine if such large-scale rearrangements were indeed homoeologous exchanges. In cases where a large rearrangement was a deletion from a NRHE, then the corresponding duplication length was defined by the common genomic positions in which the coverage showed a 1.25-fold increase compared to its mean chromosome coverage and in which the coverage increased one standard deviation from the mean. For this purpose, the coverage was calculated in 100Kb bins with bedtools *coverage* and bins with coverage above 100 were discarded to reduce mean bias due to outliers.

### Long read DNA methylation analyses

Raw reads (fast5 files) from the plants grown in Rauischholzhausen were converted from multi read to single read format using the ont_fast5_api package (Oxford Nanopore Technology) while basecalled reads were concatenated and used to annotate raw reads with tombo v.1.5.1 (Stoiber et al., 2016) using first the *annotate_raw_with_fastq* followed by the *resquiggle* functions with the *overwrite* option. Modified cytosines in CpG, CHG and CHH methylation contexts were predicted with DeepSignal-plant v. 0.1.2 (Ni et al., 2021) *call_mods* function and the *model*.*dp2*.*CNN*.*arabnrice2-1_120m_R9*.*4plus_tem*.*bn13_sn16*.*both_bilstm*.*epoch6*.*ckpt* model from the same package. The log files were then examined and only samples where the estimated coverage surpassed 30x were selected for further analyses. The frequency of methylated cytosines was calculated using the *call_freq* function and split with the *split_freq_file_by_5mC_motif*.*py* script from DeepSignal-plant. The output files were then re-merged so that they could be compatible with DMRCaller v. 1.22.0 (Catoni et al., 2018) for further differentially methylated region identification using a custom bash script (Data S1). The number of methylated cytosines and methylation level (proportion of reads supporting a methylated cytosine) in the genomic rearrangements larger than 1 Mb were calculated based on the output files and plot as heatmaps with the ComplexHeatmap (Gu et al., 2016) package. Differentially Methylated Regions (DMRs) were identified by comparing each F1 sister plant against G3D001. For this purpose, the DMRCaller *computeDMRs* function was employed to find DMRs in 1000 bp bins in chromosomes using the *bins* method, score test, a 0.01 p value threshold, and minimum cytosine count, methylation proportion difference and gap between bins of 4, 0.4 and 0 were used respectively for each variable. Lastly, DMRs were intersected with exons, introns, repeats and 1 kbp upstream promoter regions from Express 617 using bedtools.

## Results

### Large genomic rearrangements diverge across F1 sister plants

Twelve F1 sister plants derived from a single cross between Express 617 (female recipient) and G3D001 (pollen donor) were sequenced using Oxford Nanopore Technology. Sequenced reads were aligned against the Express 617 reference assembly (Lee et al., 2020). The average depth of read coverage and N50 value after read filtering were approximately 42x and 34.9 kbp, respectively (Table S1). A total of 3309 putative insertions and 1727 deletions, longer than 30 bp and supported by at least 25 reads, were detected before quality filtering as having a distinct SV allele in at least one F1 plant in comparison to the remainder of the F1 sister plants. SVs with low read allele support and low coverage were discarded. This resulted in a set of 189 and 338 high-confidence insertions and deletions respectively. PCR amplification of selected insertions from this filtered set (Table S2) confirmed that they occurred only in one or a few of the F1 sister plants (Fig. S1-S3). However, closer inspection of the sequence coverage and alleles in chromosome regions surrounding these putative SVs revealed that the detected SVs clustered in larger segmental rearrangements (Table 1) in specific chromosome regions (Table S3).

**Table 1.**
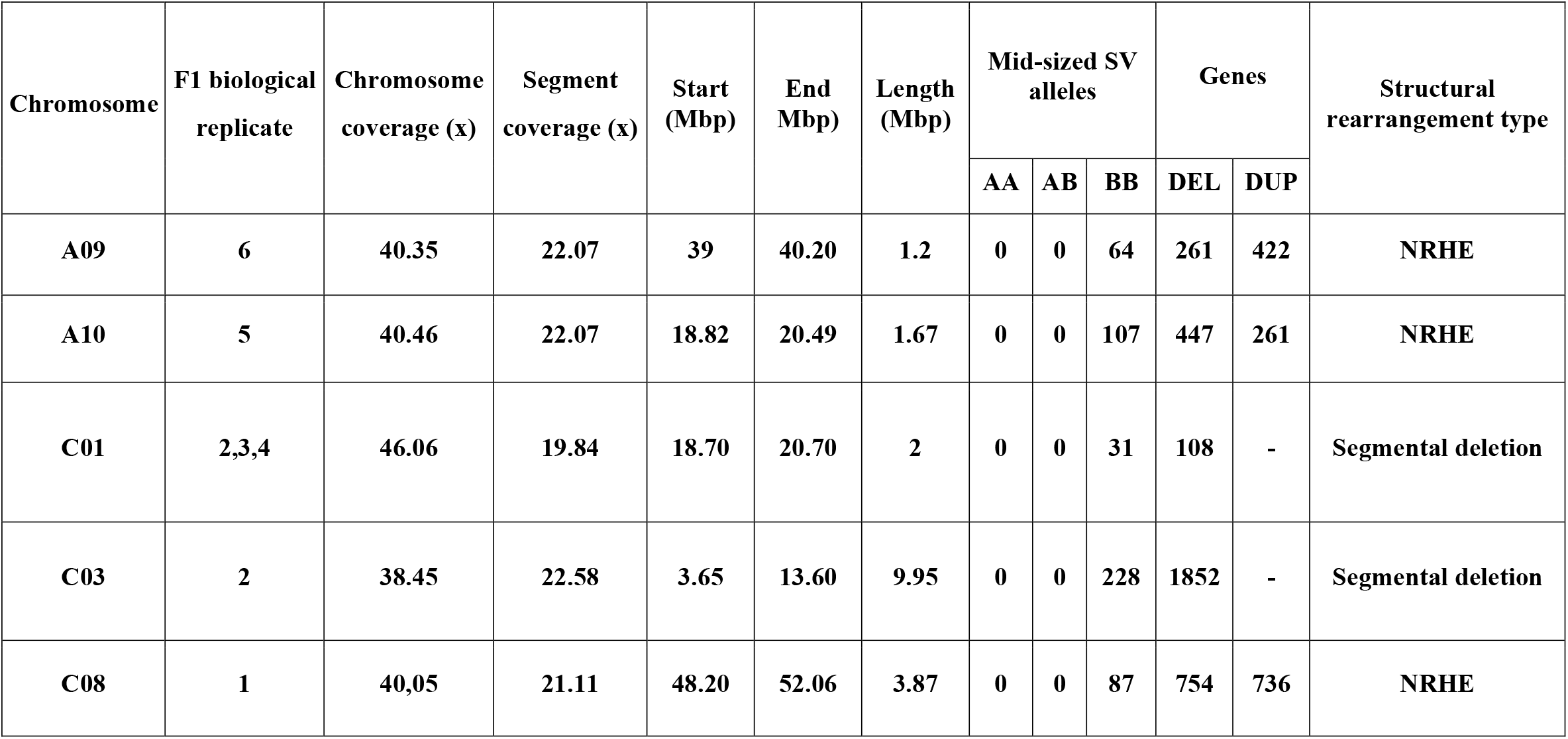
Genomic features from large segmental deletions and non-reciprocal homoeologous exchanges (NRHE). Coverage is based on the Express 617 *Brassica napus* reference genome (Lee et al., 2020). Allele information is shown as AA, AB or BB for homozygous alternate, heterozygous or homozygous reference alleles respectively. Deleted and duplicated genes within large rearrangements are displayed by the DEL and DUP abbreviations accordingly. F1 sample numbers correspond to the 12 single F1 plants.

| Chromosome | F1 biological replicate | Chromosome coverage (x) | Segment coverage (x) | Start (Mbp) | End Mbp) | Length (Mbp) | Mid-sized SV alleles |  |  | Genes |  | Structural rearrangement type |
| --- | --- | --- | --- | --- | --- | --- | --- | --- | --- | --- | --- | --- |
|  |  |  |  |  |  |  | AA | AB | BB | DEL | DUP |  |
| A09 | 6 | 40.35 | 22.07 | 39 | 40.20 | 1.2 | 0 | 0 | 64 | 261 | 422 | NRHE |
| A10 | 5 | 40.46 | 22.07 | 18.82 | 20.49 | 1.67 | 0 | 0 | 107 | 447 | 261 | NRHE |
| C01 | 2,3,4 | 46.06 | 19.84 | 18.70 | 20.70 | 2 | 0 | 0 | 31 | 108 | - | Segmental deletion |
| C03 | 2 | 38.45 | 22.58 | 3.65 | 13.60 | 9.95 | 0 | 0 | 228 | 1852 | - | Segmental deletion |
| C08 | 1 | 40,05 | 21.11 | 48.20 | 52.06 | 3.87 | 0 | 0 | 87 | 754 | 736 | NRHE |

The detected rearrangements spanned a range from 1.2 to 9.95 Mbp in length. A prominent rearrangement on chromosome C03 is displayed in Figure 1 as an example. Visualizations from large-scale rearrangements in other chromosomes are included in Figs S4-S7. As observed in Figure 1, the read coverage of chromosome C03 is mostly halved for *F1 biological replicate 2* in comparison to the rest of the genotypes and lacks insertions in this chromosome region that are specific to the paternal genotype. Interestingly, all large-scale rearrangements had a high frequency of homozygous reference alleles and halved read coverages, indicating that the segments were deleted from the inherited G3D001 chromosomes (Table 1, Figs. S4-S7) in the respective F1 individuals.

**Figure 1.**
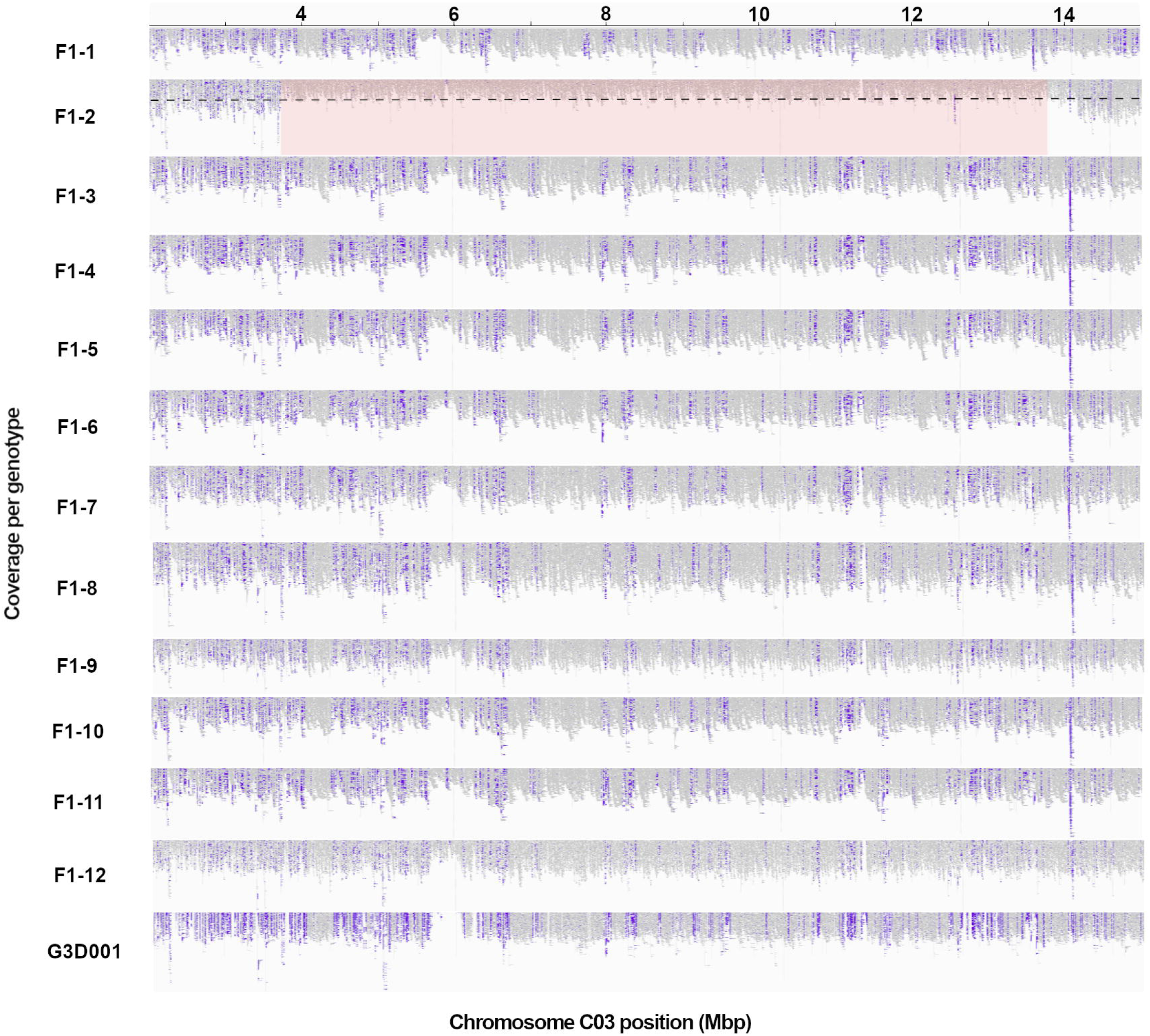
A large genomic rearrangement detected on *B. napus* chromosome C03 in a single F1 plant, *F1-2*, using Express 617 (Lee et al., 2020) as reference. Long read coverage is displayed from position 2 to 15 in Mbp using the Integrative Genomics Viewer (IGV). Purple blocks represent insertions larger than 30 bp compared to the reference assembly, as detected by IGV; as expected these are highly consistent across the 12 sister F1 plants with the exception of the segmentally deleted region in sister plant *F1-2*. The large structural rearrangement (> 1 Mbp) is highlighted in red, with a decrease in read coverage by half (as expected in a segmental deletion) represented by the black dashed line. F1 sample numbers correspond to the 12 biological replicates represented by a single F1 plant each.

Analysis of homoeologous exchanges showed that the putative segmental deletions detected in chromosomes A09, A10 and C08 in individual F1 plants are clustered in larger-scale, non-reciprocal homoeologous exchanges (Figs. S8-S10, Tables S4-S5). These NRHEs include a deleted segment from chromosome C08 that has been replaced by a duplicated segment from chromosome A09 in *F1 biological replicate 6*, a deleted segment from chromosome A10 that has been substituted by a duplicated segment from chromosome C09 in *F1 biological replicate 5*, and a deleted segment from chromosome C08 that has been replaced by a duplicated segment from chromosome A09 in *F1 biological replicate 1*.

### Impact of *de novo* SV on gene presence-absence

A total of 3422 genes were deleted and 1419 duplicated by segmental rearrangements across the 12 F1 sister offspring. Details of SV-induced gene copy number variation (CNV) are outlined in Table 1. The high rate of *de novo* genetic variation in a single, small family of F1 sister plants, reflecting the results of Higgins et al. (2018) in test-cross families, highlights the putative functional impact of chromosomal rearrangements via gene copy number variation. A clear validation of phenotype-genotype relationships is outside the scope in this study because each genotype is represented by only a single individual plant which prevents biological replicates to validate phenotypes. Nevertheless, preliminary phenotypic observations revealed large, unexpected phenological and developmental differences between individual plants. For example, 3D scanning-based phenotyping from the seedling to the full flowering stage revealed differences in plant height, leaf area and digital biomass between the *F1 biological replicate 1* and all other F1 sister plants (Figs. S11-S13, Table S6). Furthermore, this plant showed a similar phenology and development to that of Express 617, which was not the case for the other F1 sister plants. Although this might be an effect of the segmental C08 deletion and C09 duplication present in *F1 replicate 1*, nonetheless, additional F1 plants having the exact rearrangement would be required as replicates to validate the proposed hypothesis.

Additional gene copies found within the NHREs and segmental deletions in this study include *B. napus* orthologs of well-known flowering regulatory genes (*FLC, TFL1, ELF6*), along with genes corresponding to a variety of other functions (Table S7) such as disease resistance (*WRKY-4, RVB1, EDR1, EDR4, EDR8*), embryo development (*EMB1873, EMB2107, EDA22, LEA4-5*), growth and development (*DWARF4, DWARF3, OPL1, PEAR2, ATSRG1*) or abiotic stress responses (*ATHSP70-1, ATHSP90-3, ATHMP44, ATPHB2, RCI3*).

Chromosome coverage plots showed that G3D001 lacks chromosome C02 and has two copies of chromosome A02, which in turn leads to their F1 offspring having three copies of A02 and one copy of C02 (Fig. S14). The full sequences and roles in meiosis from the different A02 chromosomes in G3D001 and F1 offspring are not yet clear. Although no large-scale genomic rearrangements occurred in either of those chromosomes, further studies are still required to elucidate their impact on inheritance patterns.

### Chromosome rearrangements relate to DNA methylation patterns in F1 offspring

Genome-wide CpG, CHH, and CHG methylation was analyzed in F1 plants and their parents as described in Materials and Methods to investigate potential associations of methylation patterns with large rearrangements. The average read coverage after methylation prediction was approximately 36x (Table S8). The number of methylated cytosines was higher in the CHH context, yet the methylation level was higher in the CpG and CHG contexts (Tables S9-S10), as reported in previous studies in oilseed rape and other plants (Shen et al., 2017; Bartels et al., 2018). Overall, the number of methylated cytosines were lower in genotypes with segmental deletions as expected. Nevertheless, the methylation level was more evenly distributed among all F1 sister plants despite the presence of large segmental chromosome rearrangements (Figs. S15-S18) which could be due to uneven coverage distribution as outlined in figures (Figs. S19-S23).

The large segmental deletion on chromosome C03 (Fig. 1) was selected to illustrate methylation patterns within a large-scale structural variant. A lower number of methylated cytosines was observed in the F1 plant with the segmental deletion in the CpG methylation context in comparison to all other F1 sister plants as expected due to the deletion. Despite this, no large differences were observed in the overall methylation levels among F1 plants (Figure 2a-b). The same methylated cytosines and methylation level patterns are observed in the CHG and CHH contexts (Figure 2c-d). Differentially methylated regions were more abundant in the F1 plant with the deletion, with 36 and 75 hyper- and hypomethylated DMRs accordingly (Figure 2e). Although genes and promoters were differentially methylated, most of the genomic methylated features inside DMRs corresponded to repetitive elements (Figure 2f).

**Figure 2.**
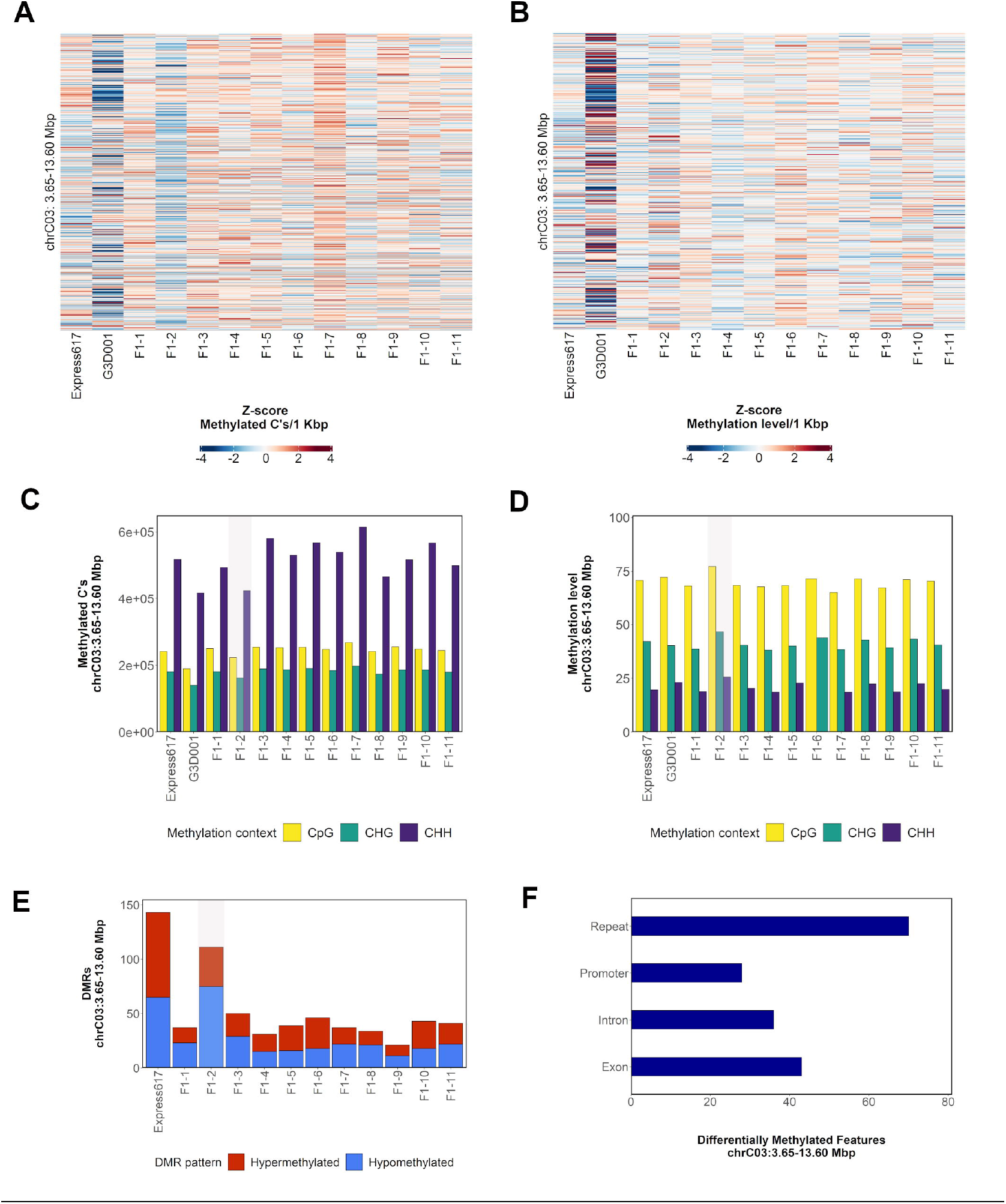
Methylation patterns in chromosome C03: 3.65– 13.60 Mbp in F1 sister plants and parents. **a**, Distribution of methylated cytosines in CpG methylation context per 1 kbp bins. Bins are sorted from bottom to top of heatmap by ascending genomic position. **b**, Distribution of methylation level in CpG methylation context per 1 kbp bins. Bins are sorted from bottom to top of heatmap by ascending genomic position. **c**, Count of methylated cytosines per methylation context. **d**, Methylation level per methylation context. **e**, Count of hypo- and hypermethylated DMRs in comparison to G3D001. **f**, Distribution of DMRs across introns, exons, repeats and promoters (1 kbp upstream from gene start). The number in the F1 samples indicates their biological replicate name. A genotype carrying a spontaneous segmental deletion on chromosome C03 is highlighted in gray.

Overall, the number of DMRs in genotypes with segmental deletions was higher, and repeats were the most prevalent methylated feature within DMRs although no repeat family enrichment was found (Figs. S15-S18). Furthermore, the average distance from DMRs to the closest gene was 2746 bp in genotypes with large-scale spontaneous rearrangements, making it feasible that they potentially play a role in transcriptomic regulation (Table S11).

In contrast to deleted chromosome segments, duplicated segments did not show any divergent methylation pattern in comparison to F1 sister plants without the corresponding duplication (Figs. S24-S29). For both deleted and duplicated regions, is noticeable that the overall methylation levels were not drastically changed. The mechanisms behind this phenomenon are still unknown.

## Discussion

A total of five large-scale, spontaneous chromosome rearrangements were observed in distinct chromosomes of different F1 sister plants. All of these rearrangements could be shown to be caused by segmental deletions occurring in inherited paternal chromosomes. The size of the rearrangements ranged from 1.2 Mbp to 9.95 Mbp and resulted not only in gene losses, but also gene duplications via non-reciprocal homoeologous exchanges. Homoeologous exchanges are known to contribute to gene loss and duplication and influence flowering time, seed lignin content and seeds per silique in *B. napus* (Stein et al., 2017; Lloyd et al., 2018). Genetic diversity within populations through other large SV such as presence-absence variation (PAV) has been previously reported in Arabidopsis, maize, sorghum and chickpea (Pucker et al., 2016; Huang et al., 2021; Tao et al., 2021; Varshney et al., 2021) as well as in oilseed rape (Gabur et al., 2018; Vollrath et al., 2021b). However, studies of PAV normally present genomic variation across genetically divergent populations of a species rather than somatic or meiotic mutations within single genotypes. Frequently, PAV analyses focus on the concept of core and disposable genes, which despite the value for pangenomic studies does not illustrate the potential regulatory role of non-coding genomic regions (Zanini et al., 2022). In this study, spontaneous exchanges and segmental deletions covering both coding and non-coding regions could be related to unexpected genetic diversity within F1 offspring from two homozygous parents.

Based on the observed parental and F1 hybrid alleles, the segmental deletions observed in the sister F1 plants most likely arose during meiosis in the pollen donor. Although they could theoretically be due to spontaneous somatic mutations, these tend to be smaller in size than the large chromosomal segments seen here. Genomic features being inherited in unexpected patterns in early generations have been reported in the form of paramutations in maize, green pea, barley grass and Arabidopsis (Lolle et al., 2005; Hollick, 2017; Adu-Yeboah et al., 2021; Bente et al., 2021; Pereira and Leitão, 2021; Cao et al., 2022), and as selfish genetic elements in rice (Lolle et al., 2005; Hollick, 2017; Yu et al., 2018). In all cases, the reported mutations were limited to a gene-size scale and not to larger genomic features. In contrast, genomic rearrangements in allotetraploid *B*.*napus* (Higgins et al. 2018) and allohexaploid Brassica hybrids (Quezada-Martinez et al. 2022) showed large scale genomic rearrangements and evidence for *de novo* SV in cross offspring that were not observed in parental lines. Given the widely reported observation of homoeologous exchanges in both synthetic and natural *B. napus* genotypes (Song et al., 1995; Szadkowski et al., 2010; Xiong et al., 2011; Samans et al., 2017; Higgins et al., 2018; Hurgobin et al., 2018), the rearrangements observed in the current study are not altogether unexpected. In contrast to previous studies, however, which used more or less densely spaced genetic markers and segregation patterns to infer positions of large-scale segmental exchanges among homoeologous chromosomes, the use of long-read sequencing enables (i) detection of SV in regions with few genetic markers, (ii) higher-resolution definition of SV breakpoints, and (iii) direct determination of the gene content and allelic composition of genes impacted by duplication and deletion events.

Species are expected to have low to intermediate mutation rates to avoid loss of required biological information (Lesaffre, 2021) and retain fitness across generations. Nevertheless, mutations within populations can lead to significant functional changes. A previous report based on sequencing of 754 plant genomes showed that annual plants carry less somatic mutations in comparison to perennials, and that the average number of mutations per biological replicate ranged from to 0.69 to 23.9 in leaf samples (Wang et al., 2019). Another comprehensive study carried out on the 25^th^ generation of a population generated by single-seed descent (SSD) in *A. thaliana* demonstrated that genomic mutations occurred randomly, and accounted for 90% of variance in gene bodies, along with accompanying epigenomic mutations (Monroe et al., 2022). In our study, a total of 3422 gene copies were deleted and 1419 were duplicated due to genomic rearrangements. Because we investigated individual, heterozygous F1 plants which cannot be biologically replicated for detailed phenotypic comparisons of seed-grown plants, a detailed analysis of phenotypic consequences from the genomic rearrangements is not possible.

In general, the individual F1 sister plants showed a very uniform phenology and morphology, as would be expected in genetically identical F1 offspring from a Mendelian cross between two largely homozygous inbred parents. However, the F1 plant *F1 replicate 1*, which was found to carry a unique NHRE between chromosomes C08 and A09 (Figs. S11-S13), was similar to the maternal line Express 617 in terms of height, digital biomass and leaf area throughout its life cycle, and dissimilar to the other 11 sister plants for these characters despite growing side-by-side in the same controlled environment. Because many genes were impacted by the various SV events, it is likely that other macro and micro-phenotypic traits could be affected by the spontaneous rearrangements in individual plants, although gene redundancy in the allopolyploid *B. napus* genome likely balances or buffers many effects from gene loss or inactivation due to rearrangements (Lesaffre, 2021). Nevertheless, the plant with a putative SV-driven impact on height, leaf area and biomass demonstrates the potential adaptive implications of frequent, spontaneous structural rearrangements as a source of novel genetic variation in a recent allopolyploid species with a narrow genetic diversity due to polyploidization and breeding bottlenecks.

Interestingly, the rearrangements on chromosomes A09, A10, C03 and C08 were located at or near telomeres, while pericentromeric regions were rearranged in chromosome C01 (Table S12). This matches corresponding observations by Higgins et al. (2018), who also observed a higher frequency of homoeologous exchanges near the ends of *B. napus* chromosomes. Distal chromosome regions tend to have a higher frequency of crossovers (Aguilar and Prieto, 2021; Kuo et al., 2021), supporting the hypothesis of Samans et al. (2017) that homoeologous rearrangements in *B. napus* are driven by meiotic crossovers between homoeologous chromosomes. This is of high relevance in a breeding context since CO occurring during meiosis results in genomic exchange, and hence, population diversity (Samans et al., 2017; Lambing and Heckmann, 2018; Mason and Wendel, 2020; Blasio et al., 2022). Although most chromosomes only exhibit one CO per meiosis in most species (Fernandes et al., 2018), it might be expected that the unusual paternal ploidy and genomic structure (Fig. S14) could have played a role in the observed F1 patterns. It has been reported that the increase or loss of specific chromosomes can alter the number of CO in *B. napus* (Suay et al., 2014).

The methylation patterns were similar to results from previous studies in oilseed rape, where methylated cytosine counts were higher in the CpG context and lower in CHH context, while methylation levels displayed the opposite trend. Likewise, DMRs were mostly found in the CpG and CHG contexts and abundantly in upstream promoter regions, as also reported previously in *B. napus* (Shen et al., 2017; Wang et al., 2018). Recent advances in long-read sequencing technology have allowed the prediction of epigenomic features like DNA cytosine methylation in Arabidopsis and triticale (Kirov et al., 2021; Naish et al., 2021). In the present study, F1 plants carrying segmental deletions displayed consistently reduced methylated cytosine counts. This is expected since the number of available cytosines that can be methylated is reduced by the deletions. Despite this, their overall methylation levels remained similar to the rest of the offspring (Fig.3, Figs. S15-S18). Furthermore, methylation levels in F1 individuals with duplicated segments was not higher than the rest of the F1 plants (Figs. S24-S26). Although DMRs were still found mostly in F1s with rearrangements, it appears that the methylation levels were maintained to similar levels across all F1 sister plants. This suggests the presence of a mechanism which maintains overall balance in methylation levels despite genomic rearrangements, for example a maternal dominance which compensates methylation losses due to deleted regions in chromosomes inherited from the paternal parent. Methylation dominance has been previously reported in resynthesized *B. napus* at the subgenome level, but the mechanisms behind this phenomenon are still not clear (Bird et al., 2021). Future work is still needed to characterize methylation patterns and the role of genomic variants and epigenomics. The implications of methylation, however, are overall key to generating diversity as it has been reported for traits such as flowering time, plant height and stress resistance (Mercé et al., 2020; Omony et al., 2020).

As in previous, related studies, we showed that genomic diversity in *B. napus* can become rapidly increased within a single generation by large scale, spontaneous chromosome rearrangements. The adaptation and survival of natural polyploids after whole genome duplication (WGD) and putative genomic shock is still not elucidated. For instance, polyploidization might lead to genomically instable offspring and reproductive isolation; however, it is also recognized as an speciation mechanism (Van de Peer et al., 2017; Pelé et al., 2018; Hörandl, 2022) and believed to contribute to environmental stress adaptation. Many species that underwent WGD have outperformed their progenitors and thrived, whereas their sister taxa did not (Van de Peer et al., 2017). Interesting examples are further described by Edger et al. (2015), who found that WGD increased genetic diversity among glucosinolate genes in Brassicales to counter herbivore predation, and by Estep et al. (2014) who discovered a considerable increase in polyploid C_4_ grasslands in the Late Miocene period. Further research is required to determine whether post-polyploidization occurs mainly through spontaneous genomic rearrangements or through environmental changes. Recent studies revealed that not all polyploidizations are linked to drastic genomic reshuffling and transcriptomic shocks, as reported in allotetraploids *A. suecica* and *B*.*rapa x Raphanus sativus* species (Burns et al., 2021; Shin et al., 2022).

The high rate of spontaneous rearrangements in the present study might lie in the synthetic nature of the paternal line, since resynthesized *B. napus* is associated with genomic instability (Szadkowski et al., 2010; Xiong et al., 2011). However, the frequency of large-scale SV is comparable to that reported by Higgins et al. (2018) in natural *B. napus*. Nevertheless, the parentage of this cross reflects potential scenarios of accelerated genomic diversity after formation of natural *B. napus*, representing an important source to enrich species diversity in a new polyploid. Our results underline previous findings showing that post-polyploidisation genome restructuring can drastically expand gene diversity among offspring in just a single self-fertilized generation. Although genetic engineering has already shown great advantages in agriculture (Sedeek et al., 2019), sudden variation generated by spontaneous chromosomes rearrangements might be an alternative method to disrupt genetic bottlenecks in scenarios where genetic engineering is not feasible.

Epigenetic modifications and structural variations altogether have contributed not only to generate diversity in the formation of allopolyploid *B. napus* (Mason and Wendel, 2020; He et al., 2021) but also in modern ecotypes (Kun Lu et al.; Song et al., 2020). Genomic rearrangements have also been associated to changes in flowering time (Schiessl et al., 2019; Chawla et al., 2021; Vollrath et al., 2021a), seed quality (Stein et al., 2017) and disease resistance (Gabur et al., 2020; Vollrath et al., 2021b) in *B. napus* cultivars. Intragenic structural variations within cultivars have also been reported in maize and wheat (Lesaffre, 2021). The present study adds a new example for rapid generation of novel genetic diversity through genome restructuring during meiosis in *B. napus*.

## Data Availability Statement

Raw and base-called reads generated and used in this study can be found in NCBI BioProject ID PRJNA837580.

## Author Contributions

RJS and JZ conceived and supervised the study. MOB drafted the manuscript and conducted the genomic and methylation bioinformatic analyses. HTL contributed to the bioinformatic pipelines and carried the centromere prediction. MM contributed to the structural variation validation, data analyses, primer design and greenhouse trials. HSC and PV contributed to the generation and interpretation of long read data and structural variations and FJS to the genomic rearrangement analysis. AL performed the 3D scanning-based phenotyping and contributed to the greenhouse trial management. RJS revised the manuscript. All authors read and approved the manuscript.

## Acknowledgements

Greenhouse and laboratory experiments were supported by Regina Illgner, Liane Renno, Stavros Tzigos, Juliette Kellermann, Birgit Keiner, Annette Plank, Daniela Quezada-Martinez and Andreas Eckert. Computational analysis was supported by the BMBF-funded de.NBI Cloud within the German Network for Bioinformatics Infrastructure (de.NBI) and the Bioinformatics Core Facility at JLU Giessen. The present research is available as a preprint at Biorxviv (Orantes-Bonilla et al., 2022).

## Supplementary Material

Supplementary Figures 1–29

Supplementary Tables 1–12

Supplementary Data 1

## References

Adu-Yeboah, P., Malone, J. M., Gill, G., and Preston, C. (2021). Non-Mendelian inheritance of gene amplification-based resistance to glyphosate in Hordeum glaucum (barley grass) from South Australia. Pest Manag Sci 77, 4298–4302. doi: 10.1002/ps.6518

Aguilar, M., and Prieto, P. (2021). Telomeres and Subtelomeres Dynamics in the Context of Early Chromosome Interactions During Meiosis and Their Implications in Plant Breeding. Front Plant Sci 12, 672489. doi: 10.3389/fpls.2021.672489

Altschul, S. F., Gish, W., Miller, W., Myers, E. W., and Lipman, D. J. (1990). Basic local alignment search tool. Journal of Molecular Biology 215, 403–410. doi: 10.1016/S0022-2836(05)80360-2

Bartels, A., Han, Q., Nair, P., Stacey, L., Gaynier, H., Mosley, M., et al. (2018). Dynamic DNA Methylation in Plant Growth and Development. Int J Mol Sci 19. doi: 10.3390/ijms19072144

Bente, H., Foerster, A. M., Lettner, N., and Mittelsten Scheid, O. (2021). Polyploidy-associated paramutation in Arabidopsis is determined by small RNAs, temperature, and allele structure. PLoS Genet 17, e1009444. doi: 10.1371/journal.pgen.1009444

Bird, K. A., Niederhuth, C. E., Ou, S., Gehan, M., Pires, J. C., Xiong, Z., et al. (2021). Replaying the evolutionary tape to investigate subgenome dominance in allopolyploid Brassica napus. New Phytol 230, 354–371. doi: 10.1111/nph.17137

Blasio, F., Prieto, P., Pradillo, M., and Naranjo, T. (2022). Genomic and Meiotic Changes Accompanying Polyploidization. Plants (Basel) 11. doi: 10.3390/plants11010125

Burns, R., Mandáková, T., Gunis, J., Soto-Jiménez, L. M., Liu, C., Lysak, M. A., et al. (2021). Gradual evolution of allopolyploidy in Arabidopsis suecica. Nat Ecol Evol 5, 1367–1381. doi: 10.1038/s41559-021-01525-w

Cai, X., and Xu, S. S. (2007). Meiosis-driven genome variation in plants. Curr Genomics 8, 151–161. doi: 10.2174/138920207780833847

Cao, S., Wang, L., Han, T., Ye, W., Liu, Y., Sun, Y., et al. (2022). Small RNAs mediate transgenerational inheritance of genome-wide trans-acting epialleles in maize. Genome Biol 23, 53. doi: 10.1186/s13059-022-02614-0

Catoni, M., Tsang, J. M., Greco, A. P., and Zabet, N. R. (2018). DMRcaller: a versatile R/Bioconductor package for detection and visualization of differentially methylated regions in CpG and non-CpG contexts. Nucleic Acids Res 46, e114. doi: 10.1093/nar/gky602

Chalhoub, B., Denoeud, F., Liu, S., Parkin, I. A. P., Tang, H., Wang, X., et al. (2014). Plant genetics. Early allopolyploid evolution in the post-Neolithic Brassica napus oilseed genome. Science 345, 950–953. doi: 10.1126/science.1253435

Chawla, H. S., Lee, H., Gabur, I., Vollrath, P., Tamilselvan-Nattar-Amutha, S., Obermeier, C., et al. (2021). Long-read sequencing reveals widespread intragenic structural variants in a recent allopolyploid crop plant. Plant Biotechnol J 19, 240–250. doi: 10.1111/pbi.13456

Cheng, C.-Y., Krishnakumar, V., Chan, A. P., Thibaud-Nissen, F., Schobel, S., and Town, C. D. (2017). Araport11: a complete reannotation of the Arabidopsis thaliana reference genome. Plant J 89, 789–804. doi: 10.1111/tpj.13415

Coster, W. de, D’Hert, S., Schultz, D. T., Cruts, M., and van Broeckhoven, C. (2018). NanoPack: visualizing and processing long-read sequencing data. Bioinformatics 34, 2666– 2669. doi: 10.1093/bioinformatics/bty149

Edger, P. P., Heidel-Fischer, H. M., Bekaert, M., Rota, J., Glöckner, G., Platts, A. E., et al. (2015). The butterfly plant arms-race escalated by gene and genome duplications. Proc Natl Acad Sci U S A 112, 8362–8366. doi: 10.1073/pnas.1503926112

Estep, M. C., McKain, M. R., Vela Diaz, D., Zhong, J., Hodge, J. G., Hodkinson, T. R., et al. (2014). Allopolyploidy, diversification, and the Miocene grassland expansion. Proc Natl Acad Sci U S A 111, 15149–15154. doi: 10.1073/pnas.1404177111

Fernandes, J. B., Séguéla-Arnaud, M., Larchevêque, C., Lloyd, A. H., and Mercier, R. (2018). Unleashing meiotic crossovers in hybrid plants. Proc Natl Acad Sci U S A 115, 2431–2436. doi: 10.1073/pnas.1713078114

Gabur, I., Chawla, H. S., Liu, X., Kumar, V., Faure, S., Tiedemann, A. von, et al. (2018). Finding invisible quantitative trait loci with missing data. Plant Biotechnol J 16, 2102–2112. doi: 10.1111/pbi.12942

Gabur, I., Chawla, H. S., Lopisso, D. T., Tiedemann, A. von, Snowdon, R. J., and Obermeier, C. (2020). Gene presence-absence variation associates with quantitative Verticillium longisporum disease resistance in Brassica napus. Sci Rep 10, 4131. doi: 10.1038/s41598-020-61228-3

Gu, Z., Eils, R., and Schlesner, M. (2016). Complex heatmaps reveal patterns and correlations in multidimensional genomic data. Bioinformatics 32, 2847–2849. doi: 10.1093/bioinformatics/btw313

Gu, Z., Gu, L., Eils, R., Schlesner, M., and Brors, B. (2014). circlize Implements and enhances circular visualization in R. Bioinformatics 30, 2811–2812. doi: 10.1093/bioinformatics/btu393

He, Z., Ji, R., Havlickova, L., Wang, L., Li, Y., Lee, H. T., et al. (2021). Genome structural evolution in Brassica crops. Nat Plants 7, 757–765. doi: 10.1038/s41477-021-00928-8

He, Z., Wang, L., Harper, A. L., Havlickova, L., Pradhan, A. K., Parkin, I. A. P., et al. (2017). Extensive homoeologous genome exchanges in allopolyploid crops revealed by mRNAseq-based visualization. Plant Biotechnol J 15, 594–604. doi: 10.1111/pbi.12657

Hickey, L. T., N Hafeez, A., Robinson, H., Jackson, S. A., Leal-Bertioli, S. C. M., Tester, M., et al. (2019). Breeding crops to feed 10 billion. Nat Biotechnol 37, 744–754. doi: 10.1038/s41587-019-0152-9

Higgins, E. E., Clarke, W. E., Howell, E. C., Armstrong, S. J., and Parkin, I. A. P. (2018). Detecting de Novo Homoeologous Recombination Events in Cultivated Brassica napus Using a Genome-Wide SNP Array. G3 (Bethesda) 8, 2673–2683. doi: 10.1534/g3.118.200118

Hohmann, M., Stahl, A., Rudloff, J., Wittkop, B., and Snowdon, R. J. (2016). Not a load of rubbish: simulated field trials in large-scale containers. Plant Cell Environ 39, 2064–2073. doi: 10.1111/pce.12737

Hollick, J. B. (2017). Paramutation and related phenomena in diverse species. Nat Rev Genet 18, 5–23. doi: 10.1038/nrg.2016.115

Hörandl, E. (2022). Novel Approaches for Species Concepts and Delimitation in Polyploids and Hybrids. Plants (Basel) 11. doi: 10.3390/plants11020204

Huang, Y., Huang, W., Meng, Z., Braz, G. T., Li, Y., Wang, K., et al. (2021). Megabase-scale presence-absence variation with Tripsacum origin was under selection during maize domestication and adaptation. Genome Biol 22, 237. doi: 10.1186/s13059-021-02448-2

Hurgobin, B., Golicz, A. A., Bayer, P. E., Chan, C.-K. K., Tirnaz, S., Dolatabadian, A., et al. (2018). Homoeologous exchange is a major cause of gene presence/absence variation in the amphidiploid Brassica napus. Plant Biotechnol J 16, 1265–1274. doi: 10.1111/pbi.12867

Jeffares, D. C., Jolly, C., Hoti, M., Speed, D., Shaw, L., Rallis, C., et al. (2017). Transient structural variations have strong effects on quantitative traits and reproductive isolation in fission yeast. Nat Commun 8, 14061. doi: 10.1038/ncomms14061

Kirov, I., Polkhovskaya, E., Dudnikov, M., Merkulov, P., Vlasova, A., Karlov, G., et al. (2021). Searching for a Needle in a Haystack: Cas9-Targeted Nanopore Sequencing and DNA Methylation Profiling of Full-Length Glutenin Genes in a Big Cereal Genome. Plants (Basel) 11. doi: 10.3390/plants11010005

Kun Lu, Lijuan Wei, Xiaolong Li, Yuntong Wang, Jian Wu, Miao Liu, et al. Whole-genome resequencing reveals Brassica napus origin and genetic loci involved in its improvement.

Kuo, P., Da Ines, O., and Lambing, C. (2021). Rewiring Meiosis for Crop Improvement. Front Plant Sci 12, 708948. doi: 10.3389/fpls.2021.708948

Labroo, M. R., Studer, A. J., and Rutkoski, J. E. (2021). Heterosis and Hybrid Crop Breeding: A Multidisciplinary Review. Front Genet 12, 643761. doi: 10.3389/fgene.2021.643761

Lambing, C., and Heckmann, S. (2018). Tackling Plant Meiosis: From Model Research to Crop Improvement. Front Plant Sci 9, 829. doi: 10.3389/fpls.2018.00829

Lee, H., Chawla, H. S., Obermeier, C., Dreyer, F., Abbadi, A., and Snowdon, R. (2020). Chromosome-Scale Assembly of Winter Oilseed Rape Brassica napus. Front Plant Sci 11, 496. doi: 10.3389/fpls.2020.00496

Lesaffre, T. (2021). Population-level consequences of inheritable somatic mutations and the evolution of mutation rates in plants. Proc Biol Sci 288, 20211127. doi: 10.1098/rspb.2021.1127

Li, H. (2018). Minimap2: pairwise alignment for nucleotide sequences. Bioinformatics 34, 3094–3100. doi: 10.1093/bioinformatics/bty191

Li, H., Handsaker, B., Wysoker, A., Fennell, T., Ruan, J., Homer, N., et al. (2009). The Sequence Alignment/Map format and SAMtools. Bioinformatics 25, 2078–2079. doi: 10.1093/bioinformatics/btp352

Lloyd, A., Blary, A., Charif, D., Charpentier, C., Tran, J., Balzergue, S., et al. (2018). Homoeologous exchanges cause extensive dosage-dependent gene expression changes in an allopolyploid crop. New Phytol 217, 367–377. doi: 10.1111/nph.14836

Lolle, S. J., Victor, J. L., Young, J. M., and Pruitt, R. E. (2005). Genome-wide non-mendelian inheritance of extra-genomic information in Arabidopsis. Nature 434, 505–509. doi: 10.1038/nature03380

Louwaars, N. P. (2018). Plant breeding and diversity: A troubled relationship? Euphytica 214, 114. doi: 10.1007/s10681-018-2192-5

Lv, H., Wang, Y., Han, F., Ji, J., Fang, Z., Zhuang, M., et al. (2020). A high-quality reference genome for cabbage obtained with SMRT reveals novel genomic features and evolutionary characteristics. Sci Rep 10, 12394. doi: 10.1038/s41598-020-69389-x

Mahmoud, M., Gobet, N., Cruz-Dávalos, D. I., Mounier, N., Dessimoz, C., and Sedlazeck, F. J. (2019). Structural variant calling: the long and the short of it. Genome Biol 20, 246. doi: 10.1186/s13059-019-1828-7

Mason, A. S., Rousseau-Gueutin, M., Morice, J., Bayer, P. E., Besharat, N., Cousin, A., et al. (2016). Centromere Locations in Brassica A and C Genomes Revealed Through Half-Tetrad Analysis. Genetics 202, 513–523. doi: 10.1534/genetics.115.183210

Mason, A.S., Higgins, E.E., Snowdon, R.J., Batley, J., Stein, A., Werner, C., et al. A user guide to the Brassica 60K Illumina Infinium™ SNP genotyping array (2017). Theor Appl Genet 130, 621–633. doi.org/10.1007/s00122-016-2849-1

Mason, A. S., and Wendel, J. F. (2020). Homoeologous Exchanges, Segmental Allopolyploidy, and Polyploid Genome Evolution. Front Genet 11, 1014. doi: 10.3389/fgene.2020.01014

Mayrose, I., Zhan, S. H., Rothfels, C. J., Arrigo, N., Barker, M. S., Rieseberg, L. H., et al. (2015). Methods for studying polyploid diversification and the dead end hypothesis: a reply to Soltis et al. (2014). New Phytol 206, 27–35. doi: 10.1111/nph.13192

Mendel, G. J. (1866). Versuche über Pflanzenhybriden. Verhandlungen des Naturforschenden Vereines in Brünn 4, 3–47.

Mercé, C., Bayer, P. E., Tay Fernandez, C., Batley, J., and Edwards, D. (2020). Induced Methylation in Plants as a Crop Improvement Tool: Progress and Perspectives. Agronomy 10, 1484. doi: 10.3390/agronomy10101484

Monroe, J. G., Srikant, T., Carbonell-Bejerano, P., Becker, C., Lensink, M., Exposito-Alonso, M., et al. (2022). Mutation bias reflects natural selection in Arabidopsis thaliana. Nature 602, 101–105. doi: 10.1038/s41586-021-04269-6

Naish, M., Alonge, M., Wlodzimierz, P., Tock, A. J., Abramson, B. W., Schmücker, A., et al. (2021). The genetic and epigenetic landscape of the Arabidopsis centromeres. Science 374, eabi7489. doi: 10.1126/science.abi7489

Ni, P., Huang, N., Nie, F., Zhang, J., Zhang, Z., Wu, B., et al. (2021). Genome-wide detection of cytosine methylations in plant from Nanopore data using deep learning. Nat Commun 12, 5976. doi: 10.1038/s41467-021-26278-9

O’Brien, K. P., Remm, M., and Sonnhammer, E. L. L. (2005). Inparanoid: a comprehensive database of eukaryotic orthologs. Nucleic Acids Res 33, D476–80. doi: 10.1093/nar/gki107

Omony, J., Nussbaumer, T., and Gutzat, R. (2020). DNA methylation analysis in plants: review of computational tools and future perspectives. Brief Bioinform 21, 906–918. doi: 10.1093/bib/bbz039

Pelé, A., Rousseau-Gueutin, M., and Chèvre, A.-M. (2018). Speciation Success of Polyploid Plants Closely Relates to the Regulation of Meiotic Recombination. Front Plant Sci 9, 907. doi: 10.3389/fpls.2018.00907

Pereira, R., and Leitão, J. M. (2021). A Non-Rogue Mutant Line Induced by ENU Mutagenesis in Paramutated Rogue Peas (Pisum sativum L.) Is Still Sensitive to the Rogue Paramutation. Genes (Basel) 12. doi: 10.3390/genes12111680

Pucker, B., Holtgräwe, D., Rosleff Sörensen, T., Stracke, R., Viehöver, P., and Weisshaar, B. (2016). A De Novo Genome Sequence Assembly of the Arabidopsis thaliana Accession Niederzenz-1 Displays Presence/Absence Variation and Strong Synteny. PLoS One 11, e0164321. doi: 10.1371/journal.pone.0164321

Quezada-Martinez, D., Zou, J., Zhang, W., Meng, J., Batley, J., and Mason, A. S. (2022). Allele segregation analysis of F1 hybrids between independent Brassica allohexaploid lineages. Chromosoma. doi: 10.1007/s00412-022-00774-3

Quinlan, A. R., and Hall, I. M. (2010). BEDTools: a flexible suite of utilities for comparing genomic features. Bioinformatics 26, 841–842. doi: 10.1093/bioinformatics/btq033

Robinson, J. T., Thorvaldsdóttir, H., Winckler, W., Guttman, M., Lander, E. S., Getz, G., et al. (2011). Integrative genomics viewer. Nat Biotechnol 29, 24–26. doi: 10.1038/nbt.1754

Rousseau-Gueutin, M., Belser, C., Da Silva, C., Richard, G., Istace, B., Cruaud, C., et al. (2020). Long-read assembly of the Brassica napus reference genome Darmor-bzh. Gigascience 9. doi: 10.1093/gigascience/giaa137

Samans, B., Chalhoub, B., and Snowdon, R. J. (2017). Surviving a Genome Collision: Genomic Signatures of Allopolyploidization in the Recent Crop Species Brassica napus. Plant Genome 10. doi: 10.3835/plantgenome2017.02.0013

Schiessl, S.-V., Katche, E., Ihien, E., Chawla, H. S., and Mason, A. S. (2019). The role of genomic structural variation in the genetic improvement of polyploid crops. The Crop Journal 7, 127–140. doi: 10.1016/j.cj.2018.07.006

Schoen, D. J., and Schultz, S. T. (2019). Somatic Mutation and Evolution in Plants. Annu. Rev. Ecol. Evol. Syst. 50, 49–73. doi: 10.1146/annurev-ecolsys-110218-024955

Sedeek, K. E. M., Mahas, A., and Mahfouz, M. (2019). Plant Genome Engineering for Targeted Improvement of Crop Traits. Front Plant Sci 10, 114. doi: 10.3389/fpls.2019.00114

Sedlazeck, F. J., Rescheneder, P., Smolka, M., Fang, H., Nattestad, M., Haeseler, A. von, et al. (2018). Accurate detection of complex structural variations using single-molecule sequencing. Nat Methods 15, 461–468. doi: 10.1038/s41592-018-0001-7

Shen, Y., Sun, S., Hua, S., Shen, E., Ye, C.-Y., Cai, D., et al. (2017). Analysis of transcriptional and epigenetic changes in hybrid vigor of allopolyploid Brassica napus uncovers key roles for small RNAs. Plant J 91, 874–893. doi: 10.1111/tpj.13605

Shin, H., Park, J. E., Park, H. R., Choi, W. L., Yu, S. H., Koh, W., et al. (2022). Admixture of divergent genomes facilitates hybridization across species in the family Brassicaceae. New Phytol. doi: 10.1111/nph.18155

Singer, S. D., Laurie, J. D., Bilichak, A., Kumar, S., and Singh, J. (2021). Genetic Variation and Unintended Risk in the Context of Old and New Breeding Techniques. Critical Reviews in Plant Sciences 40, 68–108. doi: 10.1080/07352689.2021.1883826

Soltis, D. E., Segovia-Salcedo, M. C., Jordon-Thaden, I., Majure, L., Miles, N. M., Mavrodiev, E. V., et al. (2014). Are polyploids really evolutionary dead-ends (again)? A critical reappraisal of Mayrose et al.. New Phytol 202, 1105–1117. doi: 10.1111/nph.12756

Song, J.-M., Guan, Z., Hu, J., Guo, C., Yang, Z., Wang, S., et al. (2020). Eight high-quality genomes reveal pan-genome architecture and ecotype differentiation of Brassica napus. Nat Plants 6, 34–45. doi: 10.1038/s41477-019-0577-7

Song, K., Lu, P., Tang, K., and Osborn, T. C. (1995). Rapid genome change in synthetic polyploids of Brassica and its implications for polyploid evolution. Proc Natl Acad Sci U S A 92, 7719–7723. doi: 10.1073/pnas.92.17.7719

Stein, A., Coriton, O., Rousseau-Gueutin, M., Samans, B., Schiessl, S. V., Obermeier, C., et al. (2017). Mapping of homoeologous chromosome exchanges influencing quantitative trait variation in Brassica napus. Plant Biotechnol J 15, 1478–1489. doi: 10.1111/pbi.12732

Stoiber, M., Quick, J., Egan, R., Eun Lee, J., Celniker, S., Neely, R. K., et al. (2016). De novo Identification of DNA Modifications Enabled by Genome-Guided Nanopore Signal Processing.

Suay, L., Zhang, D., Eber, F., Jouy, H., Lodé, M., Huteau, V., et al. (2014). Crossover rate between homologous chromosomes and interference are regulated by the addition of specific unpaired chromosomes in Brassica. New Phytol 201, 645–656. doi: 10.1111/nph.12534

Szadkowski, E., Eber, F., Huteau, V., Lodé, M., Huneau, C., Belcram, H., et al. (2010). The first meiosis of resynthesized Brassica napus, a genome blender. New Phytol 186, 102–112. doi: 10.1111/j.1469-8137.2010.03182.x

Tao, Y., Luo, H., Xu, J., Cruickshank, A., Zhao, X., Teng, F., et al. (2021). Extensive variation within the pan-genome of cultivated and wild sorghum. Nat Plants 7, 766–773. doi: 10.1038/s41477-021-00925-x

van de Peer, Y., Mizrachi, E., and Marchal, K. (2017). The evolutionary significance of polyploidy. Nat Rev Genet 18, 411–424. doi: 10.1038/nrg.2017.26

Varshney, R. K., Roorkiwal, M., Sun, S., Bajaj, P., Chitikineni, A., Thudi, M., et al. (2021). A chickpea genetic variation map based on the sequencing of 3,366 genomes. Nature 599, 622– 627. doi: 10.1038/s41586-021-04066-1

Vollrath, P., Chawla, H. S., Schiessl, S. V., Gabur, I., Lee, H., Snowdon, R. J., et al. (2021a). A novel deletion in FLOWERING LOCUS T modulates flowering time in winter oilseed rape. Theor Appl Genet 134, 1217–1231. doi: 10.1007/s00122-021-03768-4

Vollrath, P., Chawla, H. S., Alnajar, D., Gabur, I., Lee, H., Weber, S., et al. (2021b). Dissection of Quantitative Blackleg Resistance Reveals Novel Variants of Resistance Gene Rlm9 in Elite Brassica napus. Front Plant Sci 12, 749491. doi: 10.3389/fpls.2021.749491

Wang, L., Ji, Y., Hu, Y., Hu, H., Jia, X., Jiang, M., et al. (2019). The architecture of intra-organism mutation rate variation in plants. PLoS Biol 17, e3000191. doi: 10.1371/journal.pbio.3000191

Wang, Y., van Rengs, W. M. J., Zaidan, M. W. A. M., and Underwood, C. J. (2021). Meiosis in crops: from genes to genomes. J Exp Bot 72, 6091–6109. doi: 10.1093/jxb/erab217

Wang, Z., Wu, X., Wu, Z., An, H., Yi, B., Wen, J., et al. (2018). Genome-Wide DNA Methylation Comparison between Brassica napus Genic Male Sterile Line and Restorer Line. Int J Mol Sci 19. doi: 10.3390/ijms19092689

Xiong, Z., Gaeta, R. T., and Pires, J. C. (2011). Homoeologous shuffling and chromosome compensation maintain genome balance in resynthesized allopolyploid Brassica napus. Proc Natl Acad Sci U S A 108, 7908–7913. doi: 10.1073/pnas.1014138108

Yu, X., Zhao, Z., Zheng, X., Zhou, J., Kong, W., Wang, P., et al. (2018). A selfish genetic element confers non-Mendelian inheritance in rice. Science 360, 1130–1132. doi: 10.1126/science.aar4279

Yuan, Y., Bayer, P. E., Batley, J., and Edwards, D. (2021). Current status of structural variation studies in plants. Plant Biotechnol J 19, 2153–2163. doi: 10.1111/pbi.13646

Zanini, S. F., Bayer, P. E., Wells, R., Snowdon, R. J., Batley, J., Varshney, R. K., et al. (2022). Pangenomics in crop improvement-from coding structural variations to finding regulatory variants with pangenome graphs. Plant Genome 15, e20177. doi: 10.1002/tpg2.20177

Zhang, L., Cai, X., Wu, J., Liu, M., Grob, S., Cheng, F., et al. (2018). Improved Brassica rapa reference genome by single-molecule sequencing and chromosome conformation capture technologies. Hortic Res 5, 50. doi: 10.1038/s41438-018-0071-9

Zou, J., Hu, D., Mason, A. S., Shen, X., Wang, X., Wang, N., et al. (2018). Genetic changes in a novel breeding population of Brassica napus synthesized from hundreds of crosses between B. rapa and B. carinata. Plant Biotechnol J 16, 507–519. doi: 10.1111/pbi.12791

