## Supplementary Figures for "Frequent spontaneous structural rearrangements promote rapid genome diversification in a *Brassica napus* F1 generation"

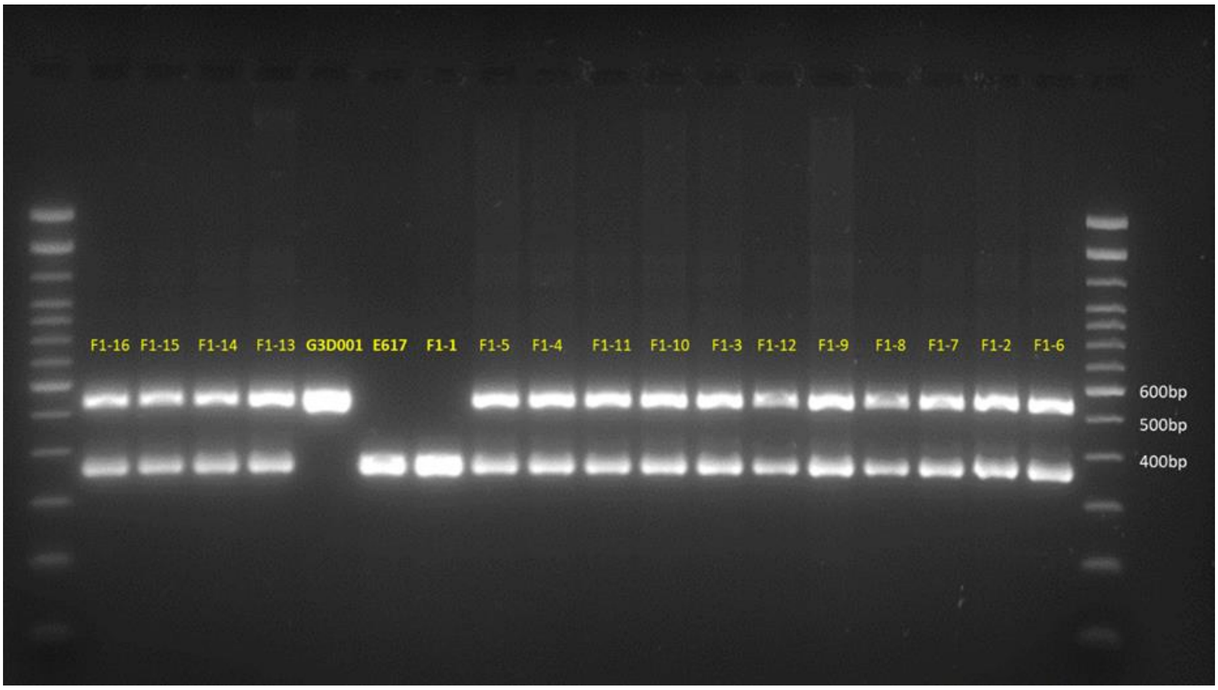


**Supplementary Figure 1. Gel electrophoresis image from a 115 bp insertion in chromosome C08.** Reference fragment sizes are on white labels and replicate names are on yellow. E617 and G3D001 corresponds to the Express 617 and G3D001 biological replicates crossed to develop all F1 sister plants respectively, whereas the number in the F1 samples indicates their biological replicate name.

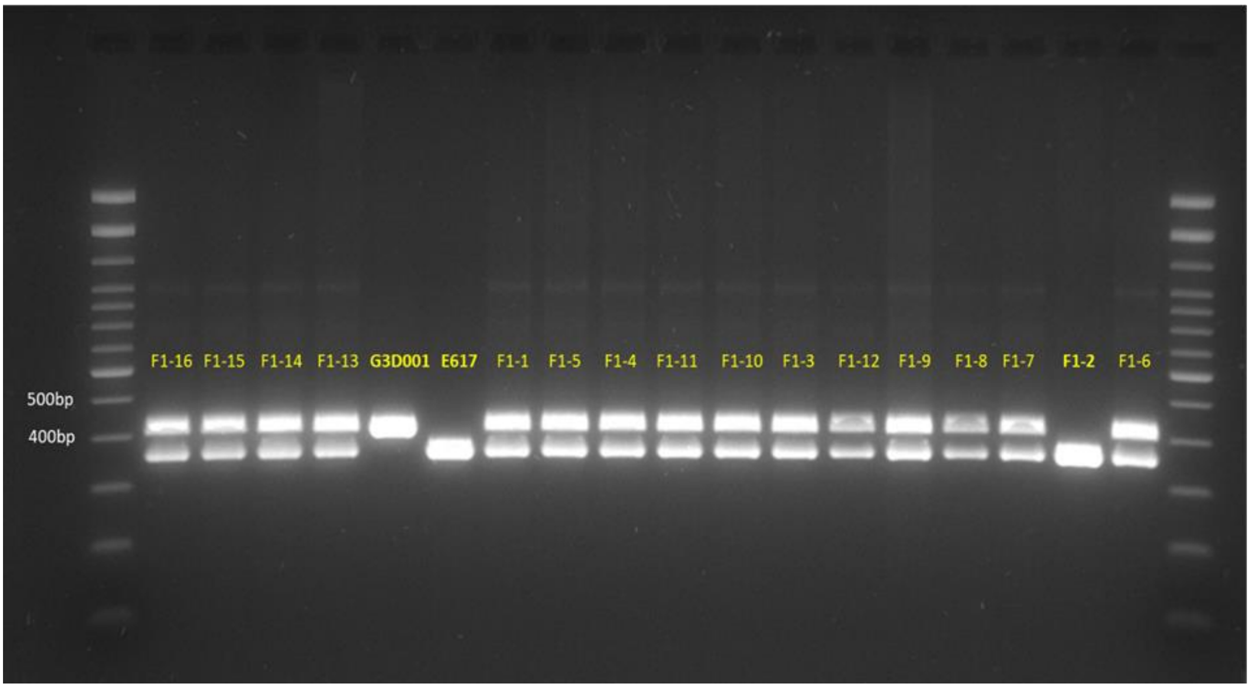


**Supplementary Figure 2. Gel electrophoresis image from a 60 bp insertion in chromosome C03.** Reference fragment sizes are on white labels and replicate names are on yellow. E617 and G3D001 corresponds to the Express 617 and G3D001 biological replicates crossed to develop all F1 sister plants, whereas the number in the F1 samples indicates their biological replicate name.

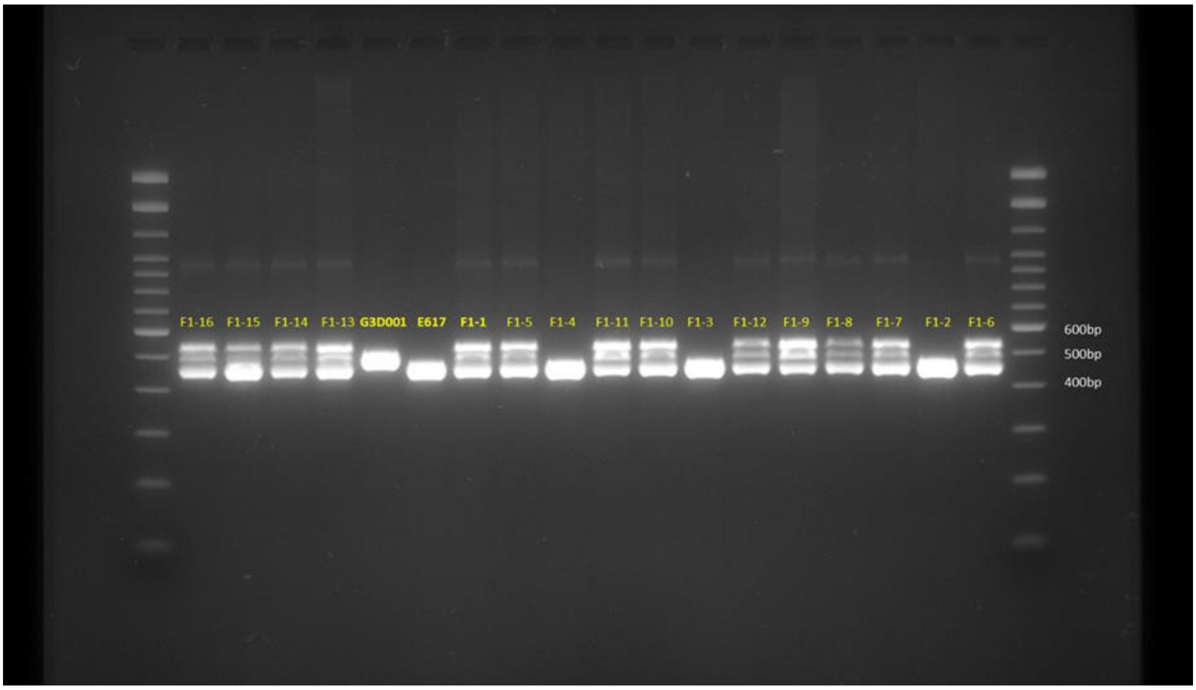


**Supplementary Figure 3. Gel electrophoresis image from a 37 bp insertion in chromosome C01.** Reference fragment sizes are on white labels and replicate names are on yellow. E617 and G3D001 corresponds to the Express 617 and G3D001 biological replicates crossed to develop all F1 sister plants, whereas the number in the F1 samples indicates their biological replicate name.

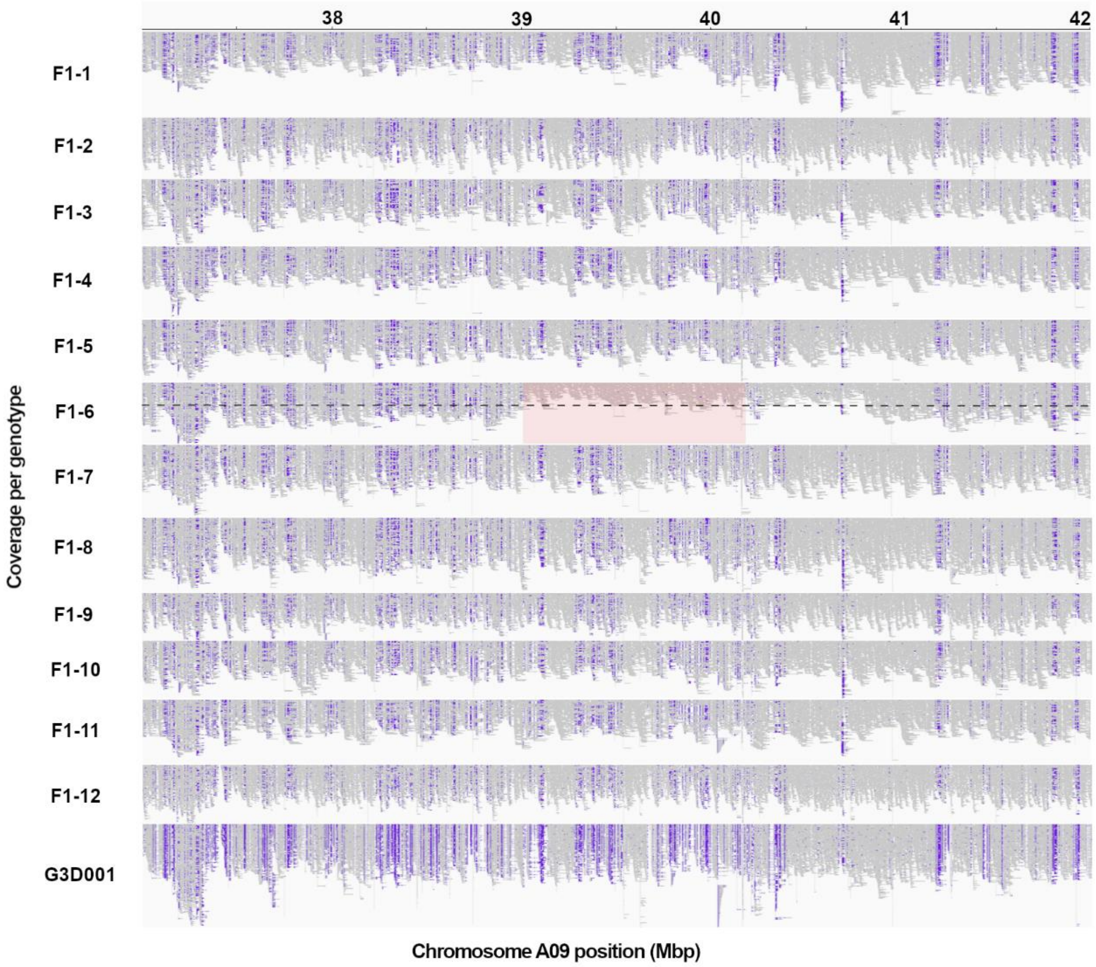


**Supplementary Figure 4. A large genomic rearrangement detected on *B. napus* chromosome A09 in a single F1 plant, *F1-6*, using Express 617 (Lee et al., 2020) as reference.** Long read coverage is displayed from position 37 to 42 in Mbp using the Integrative Genomics Viewer (IGV). Purple blocks represent insertions larger than 30 bp compared to the reference assembly, as detected by IGV. The large structural rearrangement (> 1 Mbp) is highlighted in red, with a decrease in read coverage by half represented by the black dashed line. F1 sample numbers correspond to the 12 biological replicates represented by a single F1 plant each.

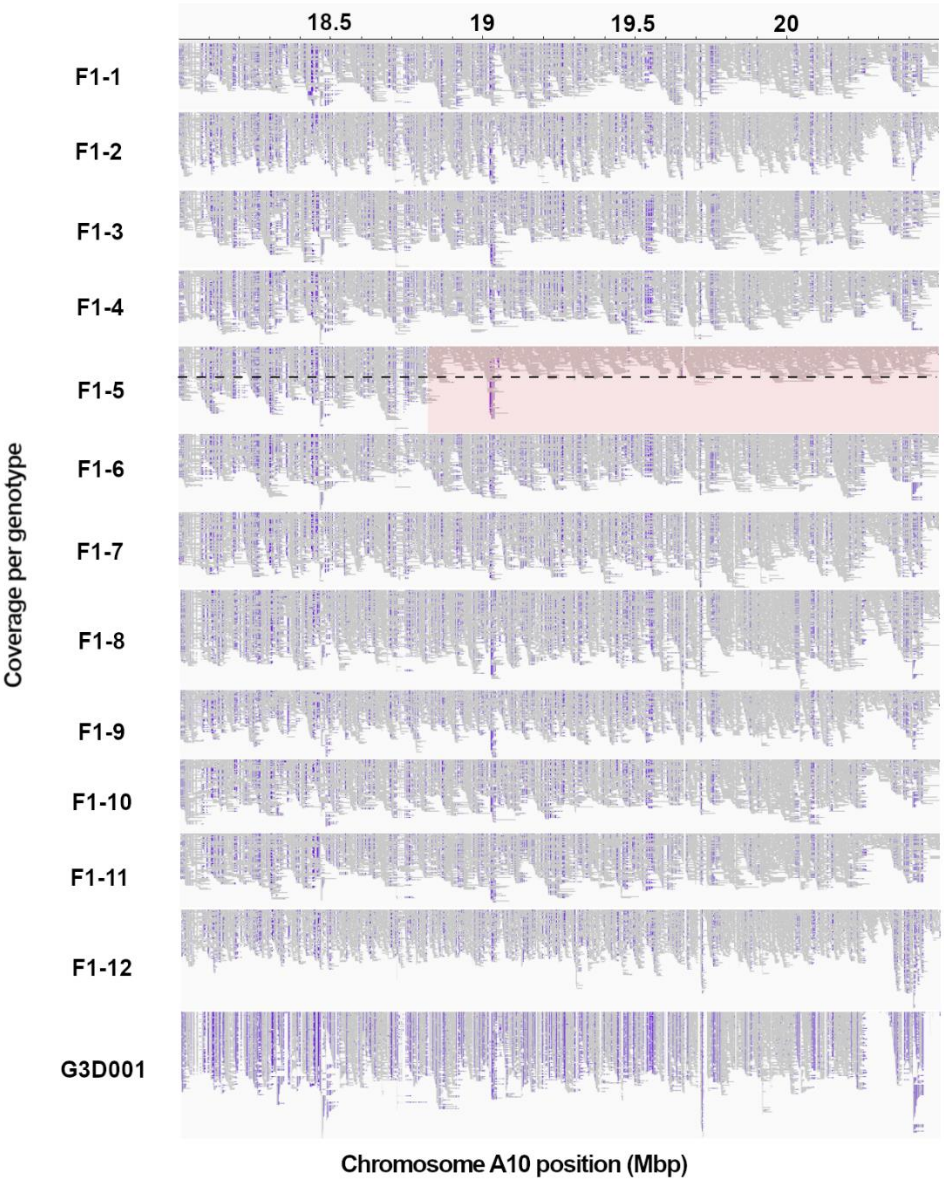


**Supplementary Figure 5. A large genomic rearrangement detected on *B. napus* chromosome A10 in a single F1 plant, *F1-5*, using Express 617 (Lee et al., 2020) as reference.** Long read coverage is displayed from position 18 to 20.49 in Mbp using the Integrative Genomics Viewer (IGV). Purple blocks represent insertions larger than 30 bp compared to the reference assembly, as detected by IGV. The large structural rearrangement (> 1 Mbp) is highlighted in red, with a decrease in read coverage by half represented by the black dashed line. F1 sample numbers correspond to the 12 biological replicates represented by a single F1 plant each.

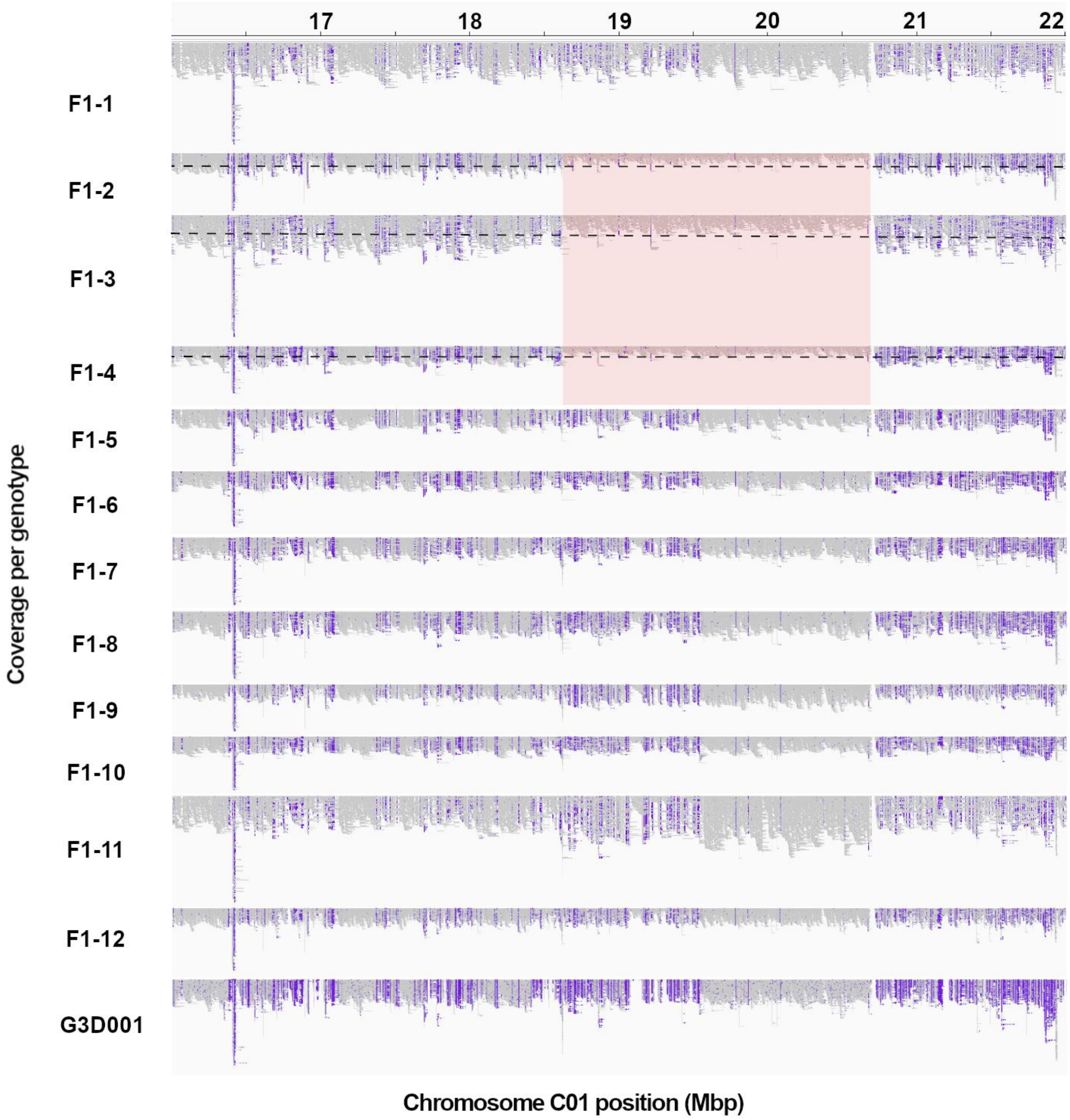


**Supplementary Figure 6. A large genomic rearrangement detected on *B. napus* chromosome C01 in a three F1 plants, *F1-2*, *F1-3* and *F1-4*, using Express 617 (Lee et al., 2020) as reference.** Long read coverage is displayed from position 16 to 22 in Mbp using the Integrative Genomics Viewer (IGV). Purple blocks represent insertions larger than 30 bp compared to the reference assembly, as detected by IGV. The large structural rearrangement (> 1 Mbp) is highlighted in red, with a decrease in read coverage by half represented by the black dashed line. F1 sample numbers correspond to the 12 biological replicates represented by a single F1 plant each.

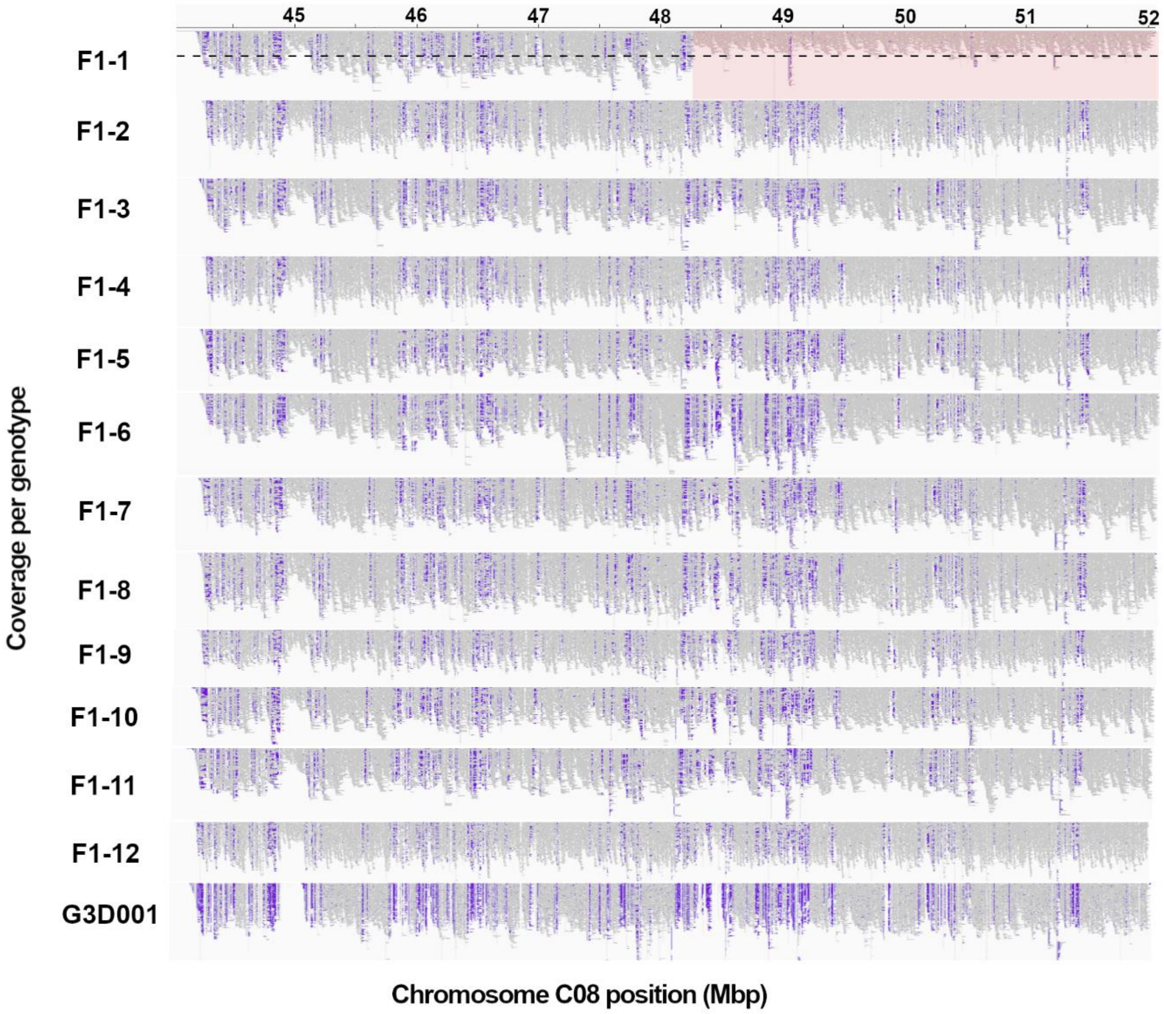


**Supplementary Figure 7. A large genomic rearrangement detected on *B. napus* chromosome C08 in a single F1 plant, *F1-1*, using Express 617 (Lee et al., 2020) as reference.** Long read coverage is displayed from position 44 to 52.06 in Mbp using the Integrative Genomics Viewer (IGV). Purple blocks represent insertions larger than 30 bp compared to the reference assembly, as detected by IGV. The large structural rearrangement (> 1 Mbp) is highlighted in red, with a decrease in read coverage by half represented by the black dashed line. F1 sample numbers correspond to the 12 biological replicates represented by a single F1 plant each.

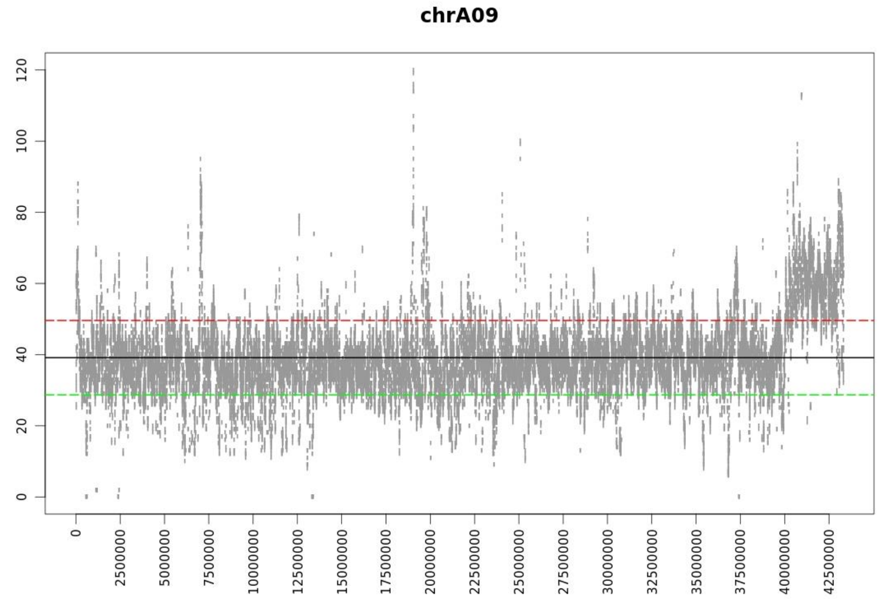


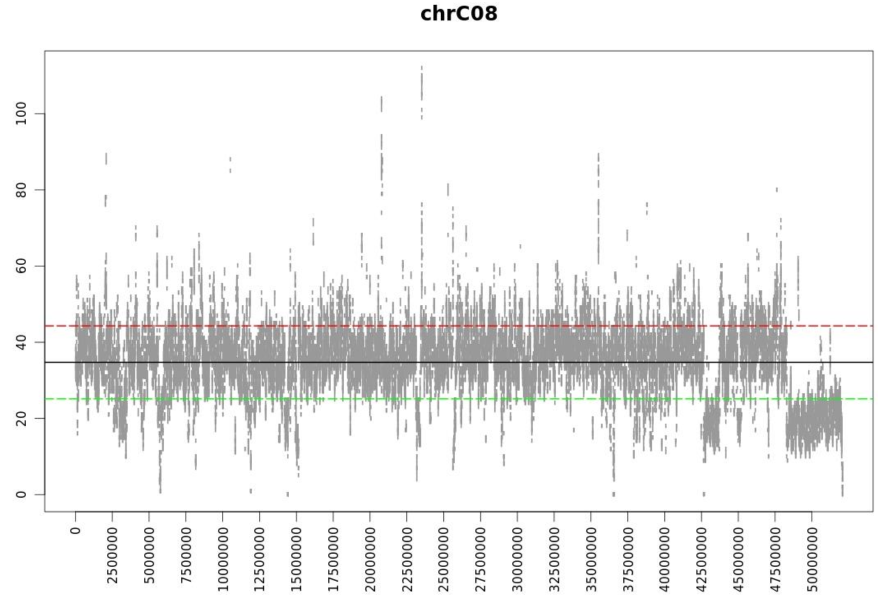


**Supplementary Figure 8. Coverage of chromosome C08 and homoeologous chromosome A09 in the Express617xG3D001 F1 biological replicate 1 based on the Express 617 (Lee et al., 2020) reference.** Duplications are shown above the red line and deletions below the green line.

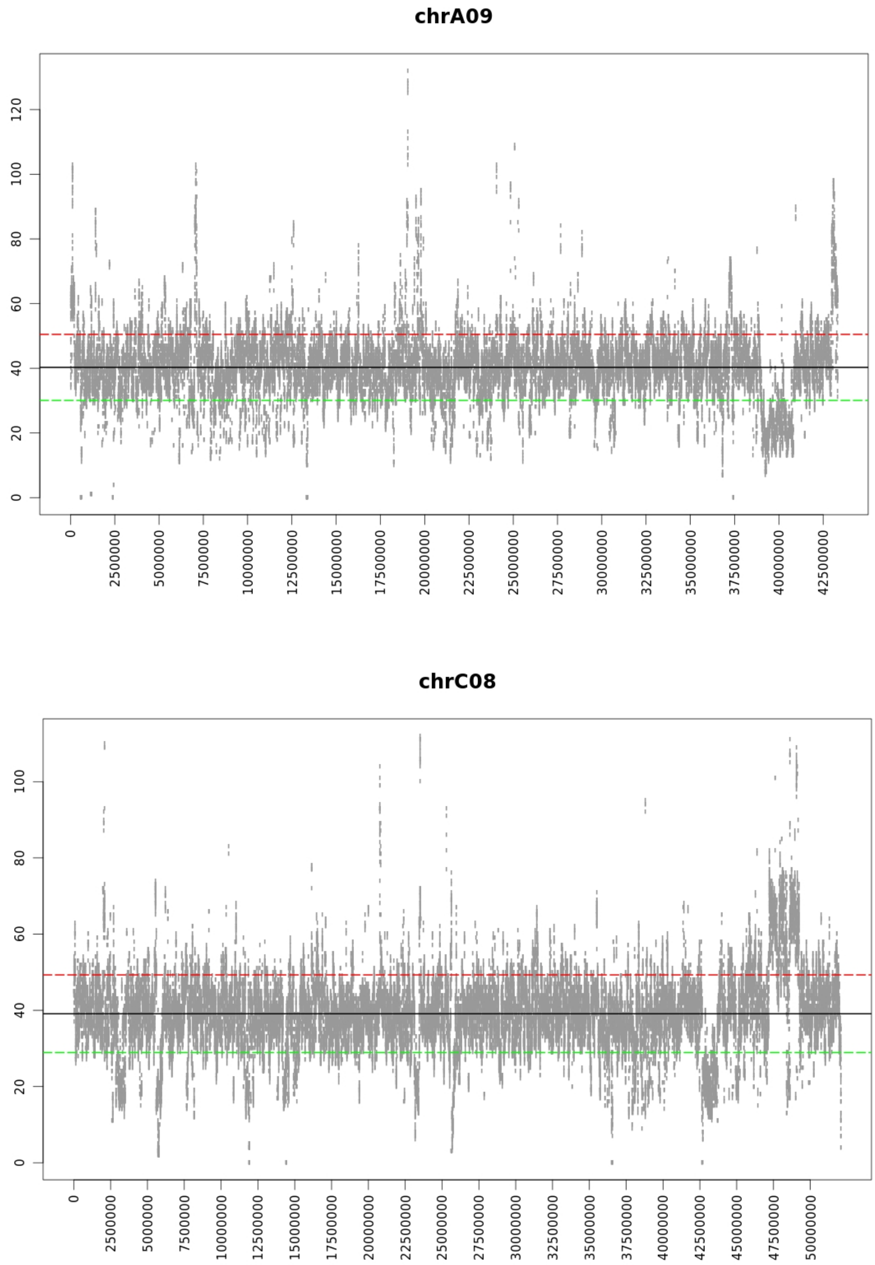


**Supplementary Figure 9. Coverage of chromosome A09 and homoeologous chromosome C08 in the Express617xG3D001 F1 biological replicate 6 based on the Express 617 (Lee et al., 2020) reference.** Duplications are shown above the red line and deletions below the green line.

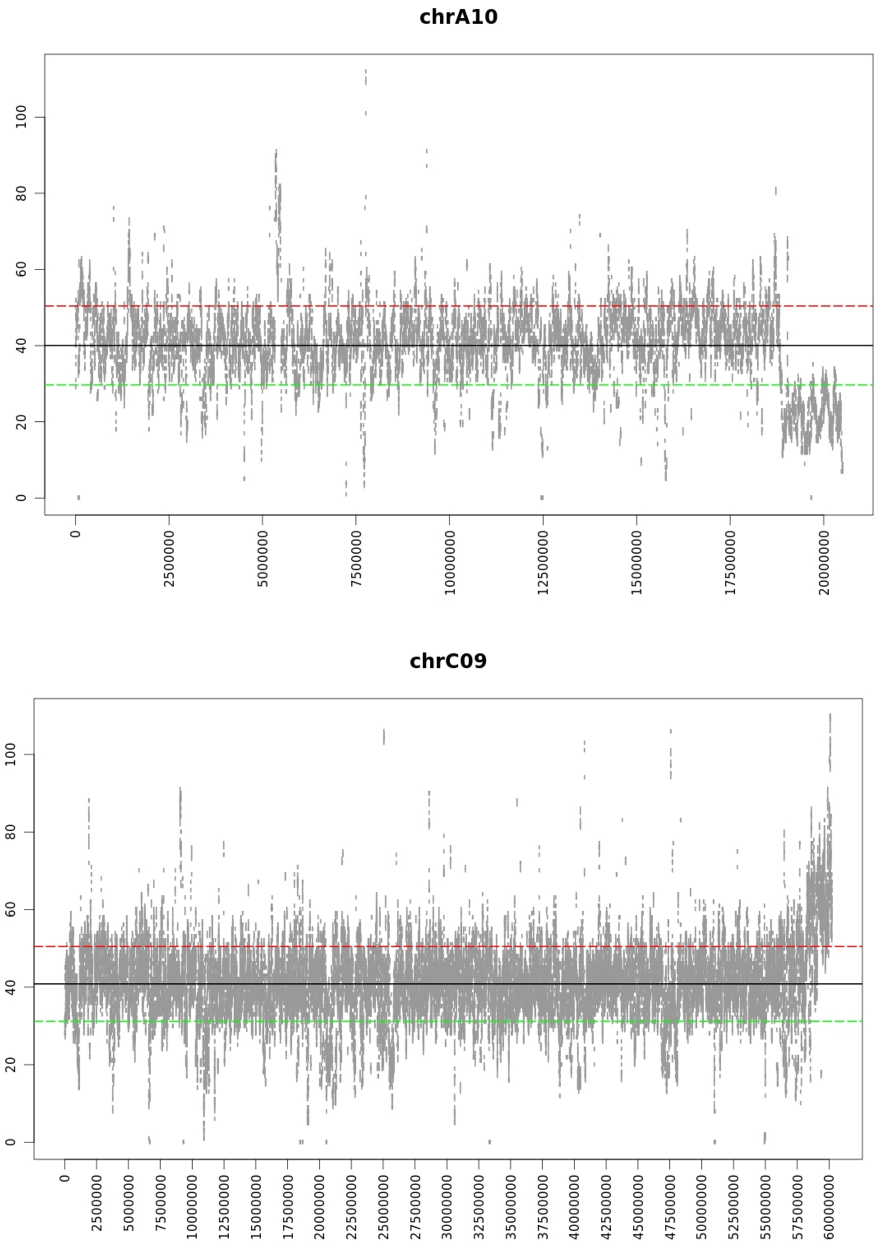


**Supplementary Figure 10. Coverage of chromosome A10 and homoeologous chromosome C09 in the Express617xG3D001 F1 biological replicate 5 based on the Express 617 (Lee et al., 2020) reference.** Duplications are shown above the red line and deletions below the green line.

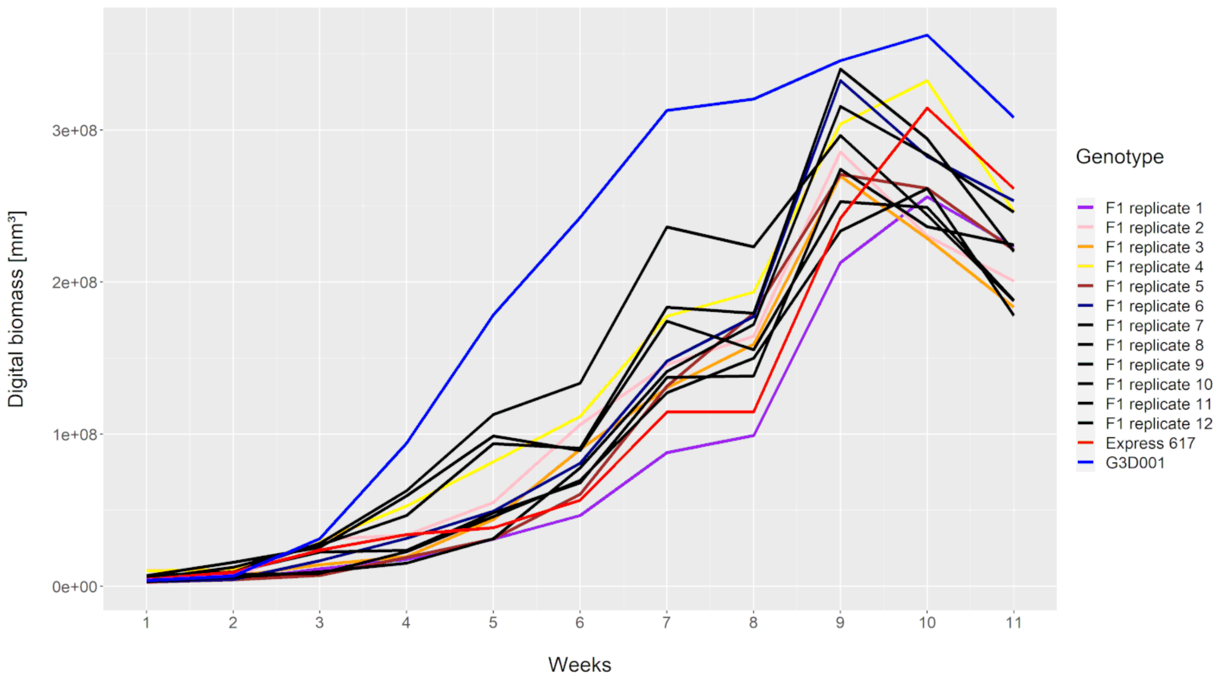


**Supplementary Figure 11. Digital biomass ($mm^3$) records per genotype from seedling stage to full flowering.** Only the parents and F1 hybrids with large rearrangements (> 1Mbp) are displayed in colors, whereas genotypes without large rearrangements are shown in black.

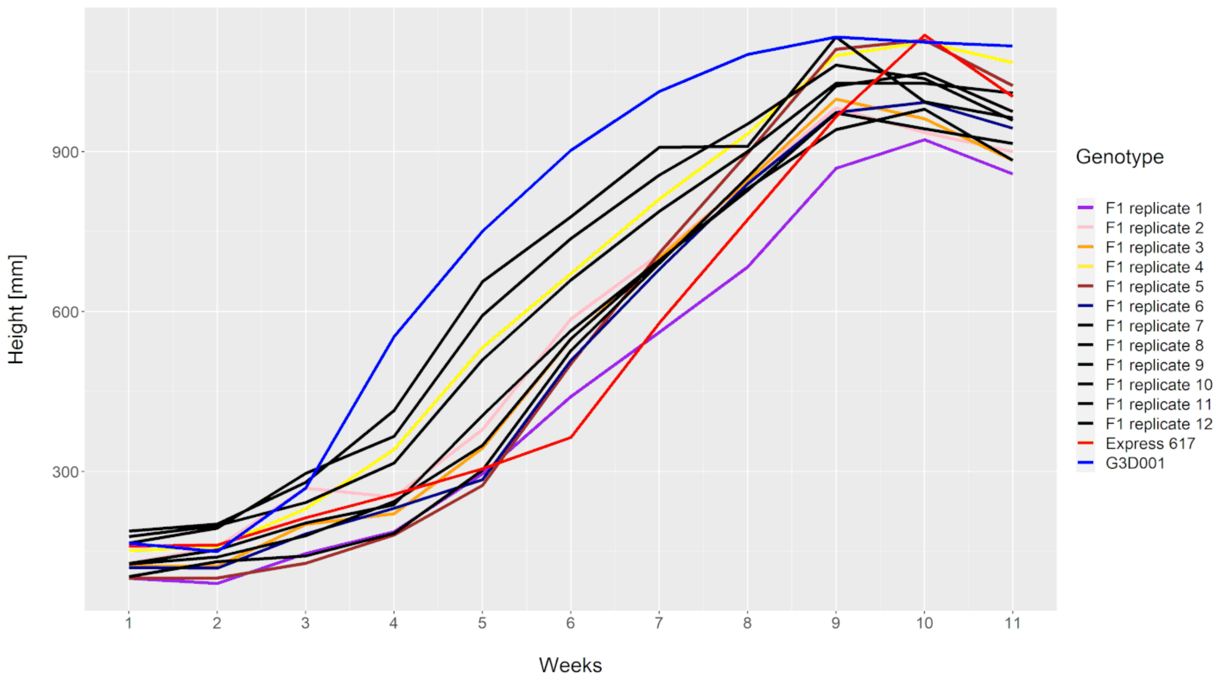


**Supplementary Figure 12. Height (mm) records per genotype from seedling stage to full flowering.** Only the parents and F1 sister plants with large rearrangements (> 1Mbp) are displayed in colors, whereas genotypes without large rearrangements are shown in black.

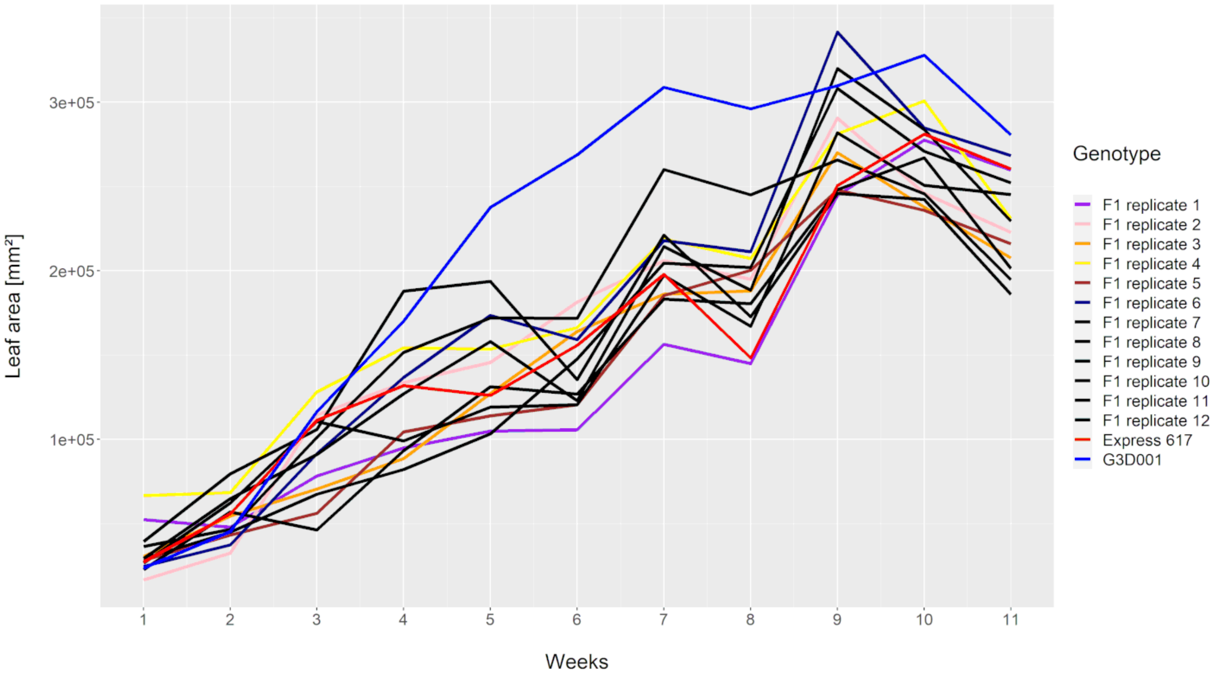


**Supplementary Figure 13. Leaf area (mm$^3$) records per genotype from seedling stage to full flowering.** Only the parents and F1 sister plants with large rearrangements (> 1Mbp) are displayed in colors, whereas genotypes without large rearrangements are shown in black.

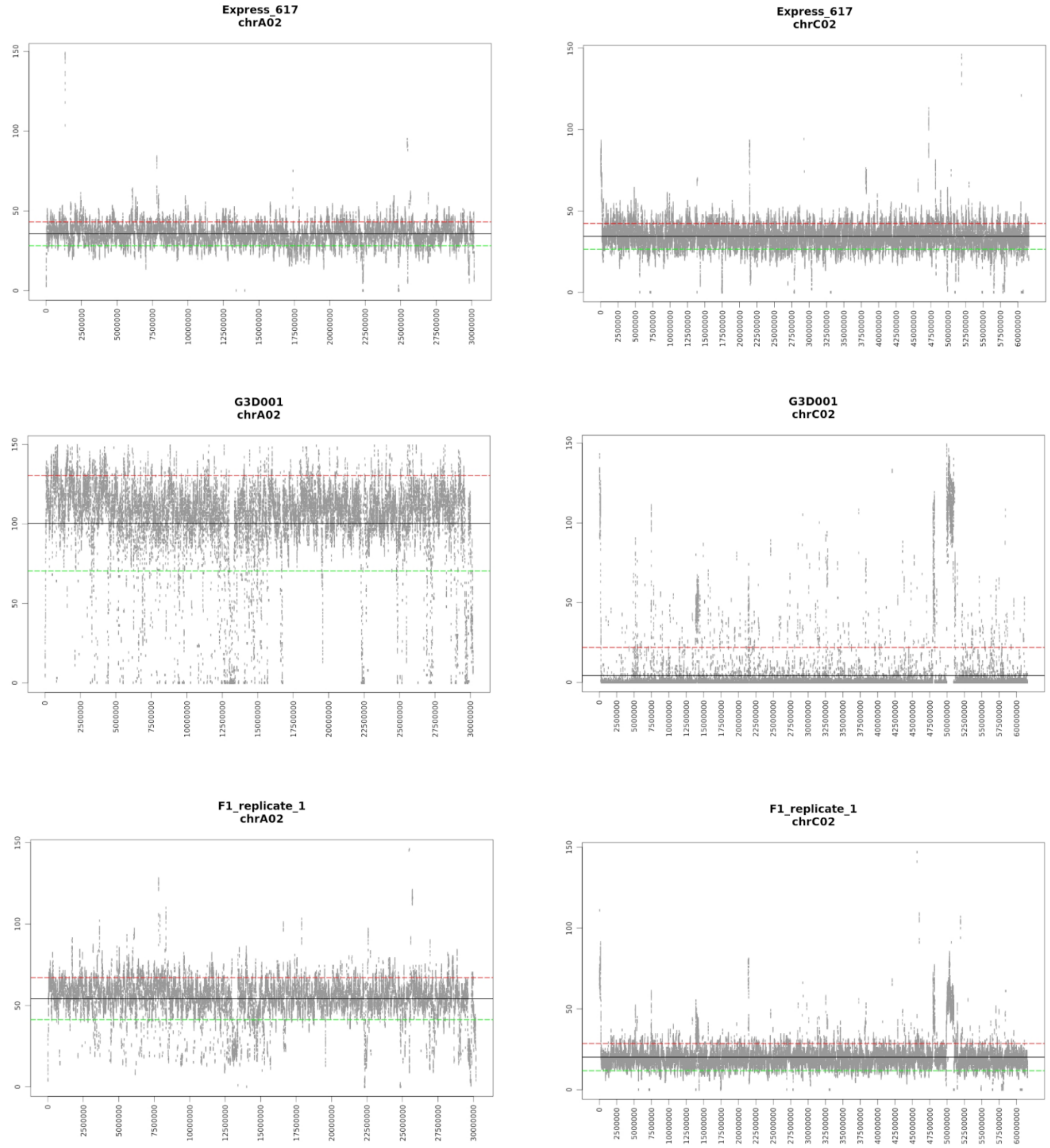


**Supplementary Figure 14. Coverage of chromosomes A02 and C02 from Express 617, G3D001 and selected F1 based on the Express 617 (Lee et al., 2020) reference.** Duplications are shown above the red line and deletions below the green line.

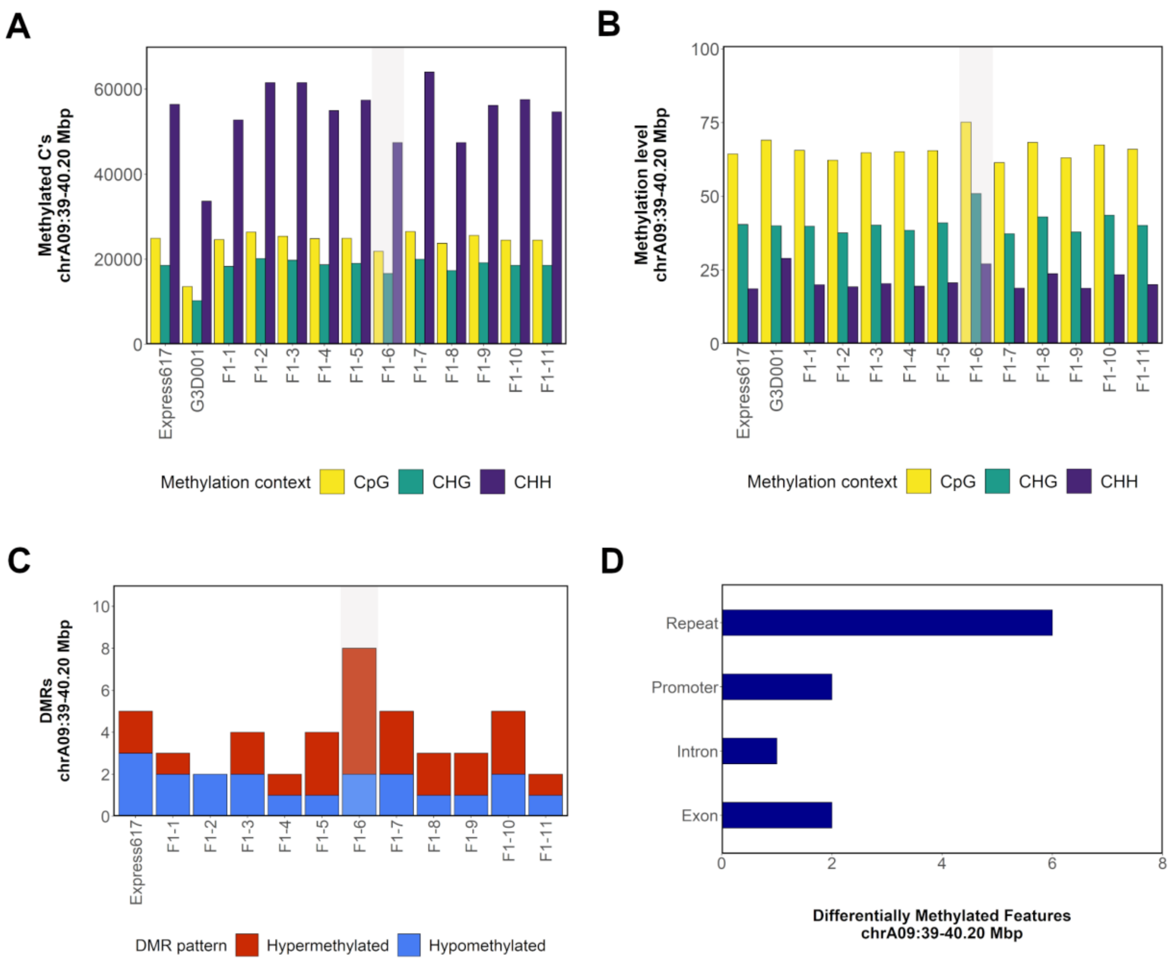


**Supplementary Figure 15. Methylation patterns in chromosome A09: 39– 40.2 Mbp. a**, Count of methylated cytosines per methylation context. **b**, Methylation level per methylation context. **c**, Count of hypo- and hypermethylated DMRs in comparison to G3D001. **d**, Distribution of DMRs across introns, exons, repeats and promoters (1 kbp upstream from gene start). The number in the F1 samples indicates their biological replicate name. A genotype carrying a spontaneous NRHE on chromosome A09 is highlighted in gray.

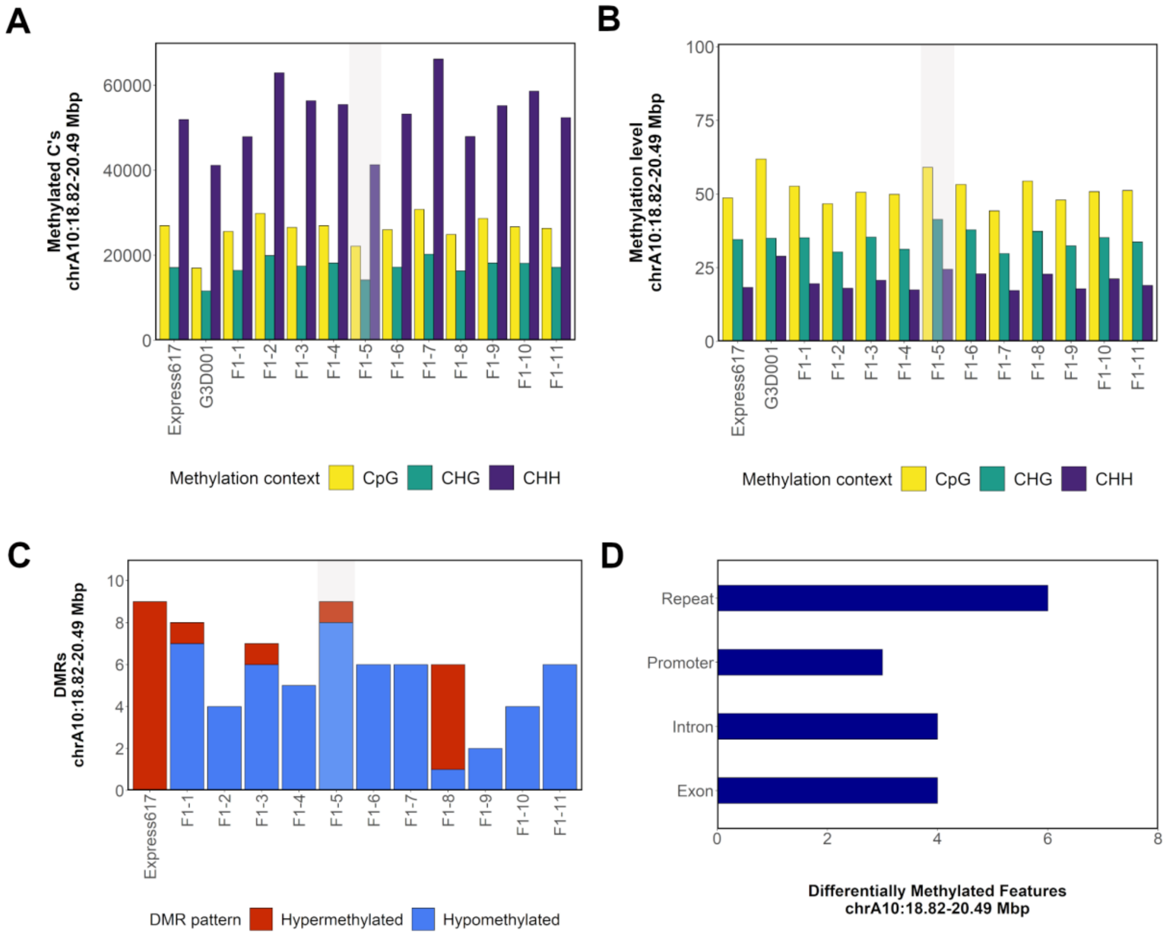


**Supplementary Figure 16. Methylation patterns in chromosome A10: 18.82– 20.49 Mbp.** **a**, Count of methylated cytosines per methylation context. **b**, Methylation level per methylation context. **c**, Count of hypo- and hypermethylated DMRs in comparison to G3D001. **d**, Distribution of DMRs across introns, exons, repeats and promoters (1 kbp upstream from gene start). The number in the F1 samples indicates their biological replicate name. A genotype carrying a spontaneous NRHE on chromosome A10 is highlighted in gray.

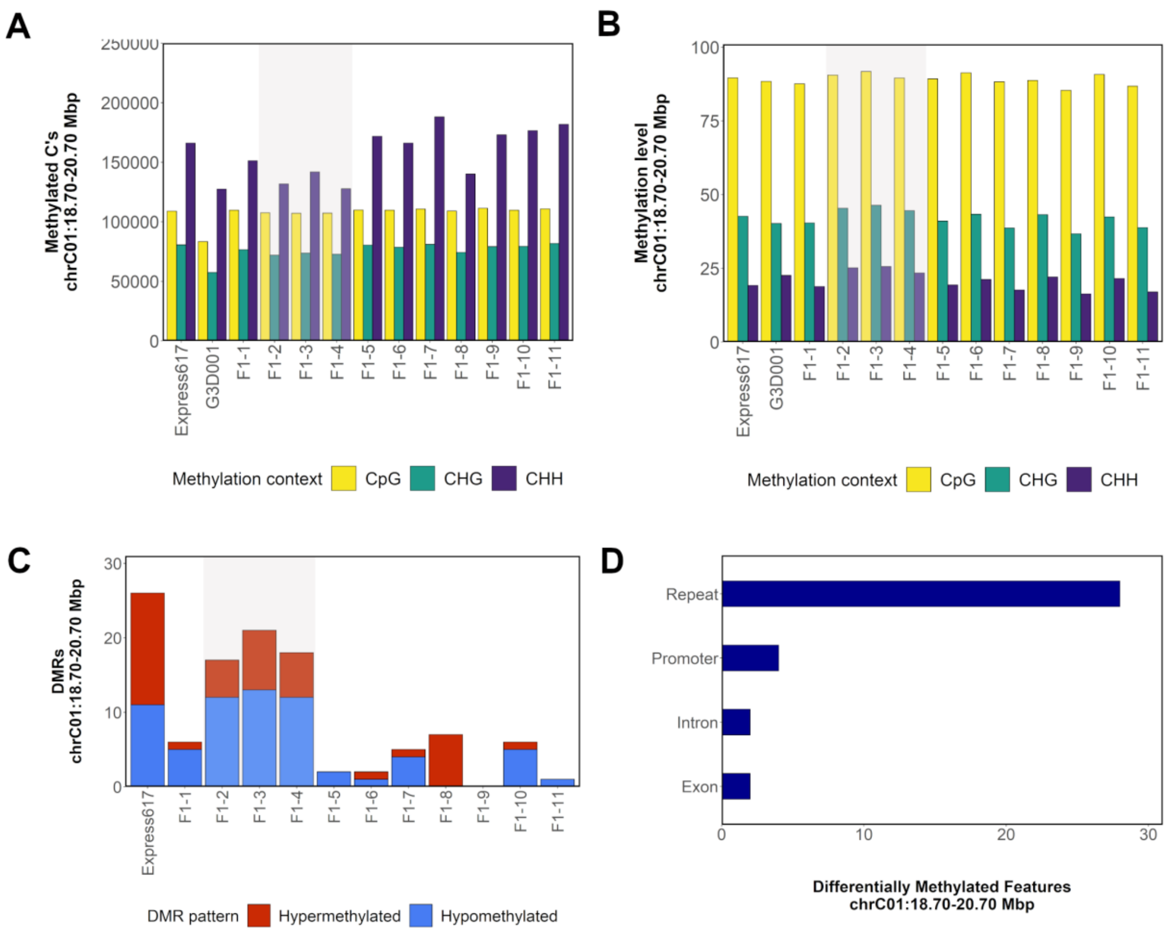


**Supplementary Figure 17. Methylation patterns in chromosome C01: 18.7– 20.7 Mbp. a**, Count of methylated cytosines per methylation context. **b**, Methylation level per methylation context. **c**, Count of hypo- and hypermethylated DMRs in comparison to G3D001. **d**, Distribution of DMRs across introns, exons, repeats and promoters (1 kbp upstream from gene start). The number in the F1 samples indicates their biological replicate name. Genotypes carrying a spontaneous segmental deletion on chromosome C01 are highlighted in gray.

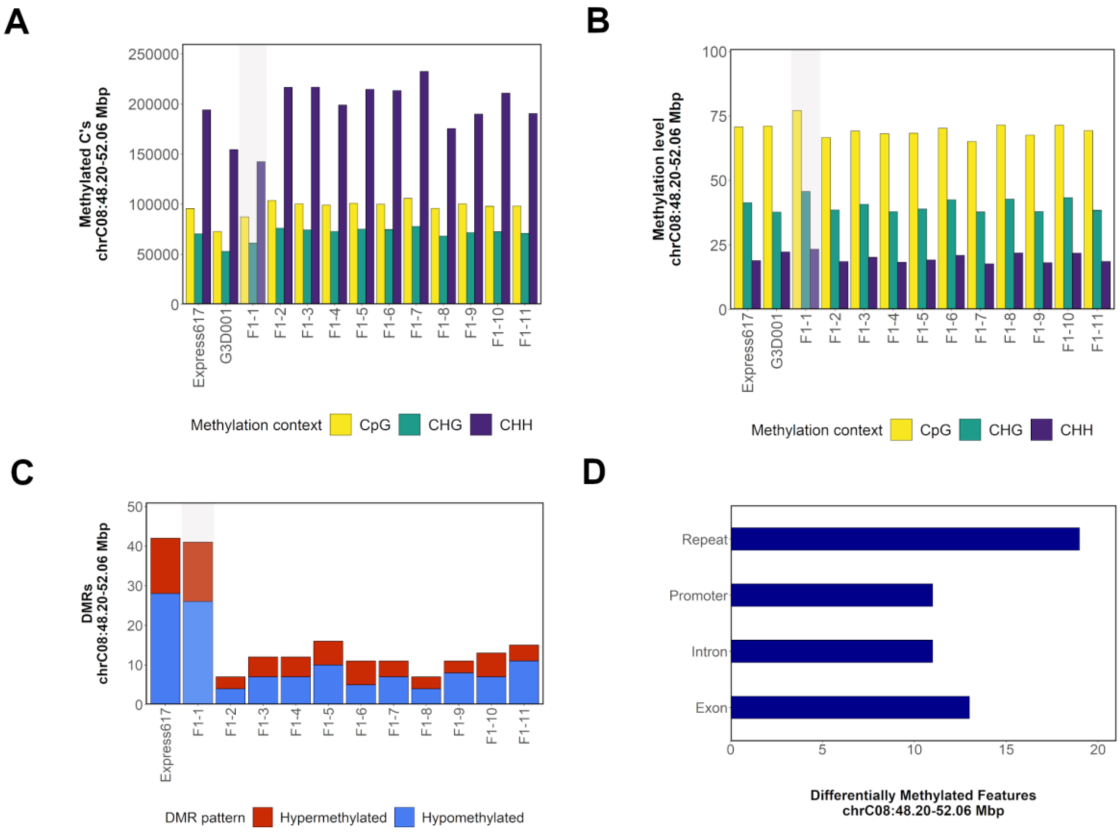


**Supplementary Figure 18. Methylation patterns in chromosome C08: 3.65– 13.60. a**, Count of methylated cytosines per methylation context. **b**, Methylation level per methylation context. **c**, Count of hypo- and hypermethylated DMRs in comparison to G3D001. **d**, Distribution of DMRs across introns, exons, repeats and promoters (1 kbp upstream from gene start). The number in the F1 samples indicates their biological replicate name. A genotype carrying a spontaneous NRHE on chromosome C08 is highlighted in gray.

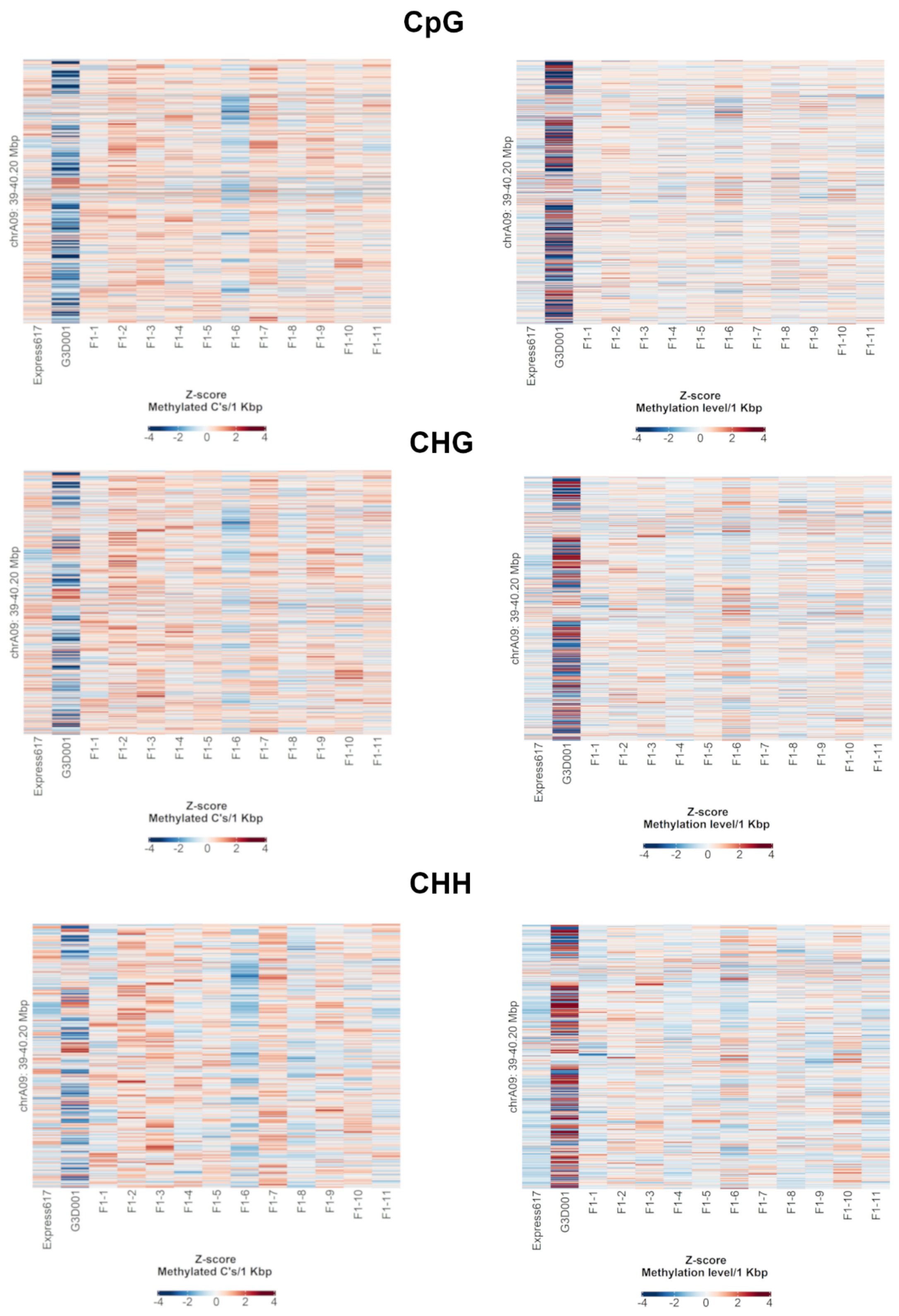


**Supplementary Figure 19. Count of methylated cytosines (left) and methylation level (right) per methylation context within 1 kbp bins across F1 sister plants and parents in chromosome A09: 39– 40.2 Mbp.** Bins are sorted from bottom to top from heatmaps by ascending genomic position.

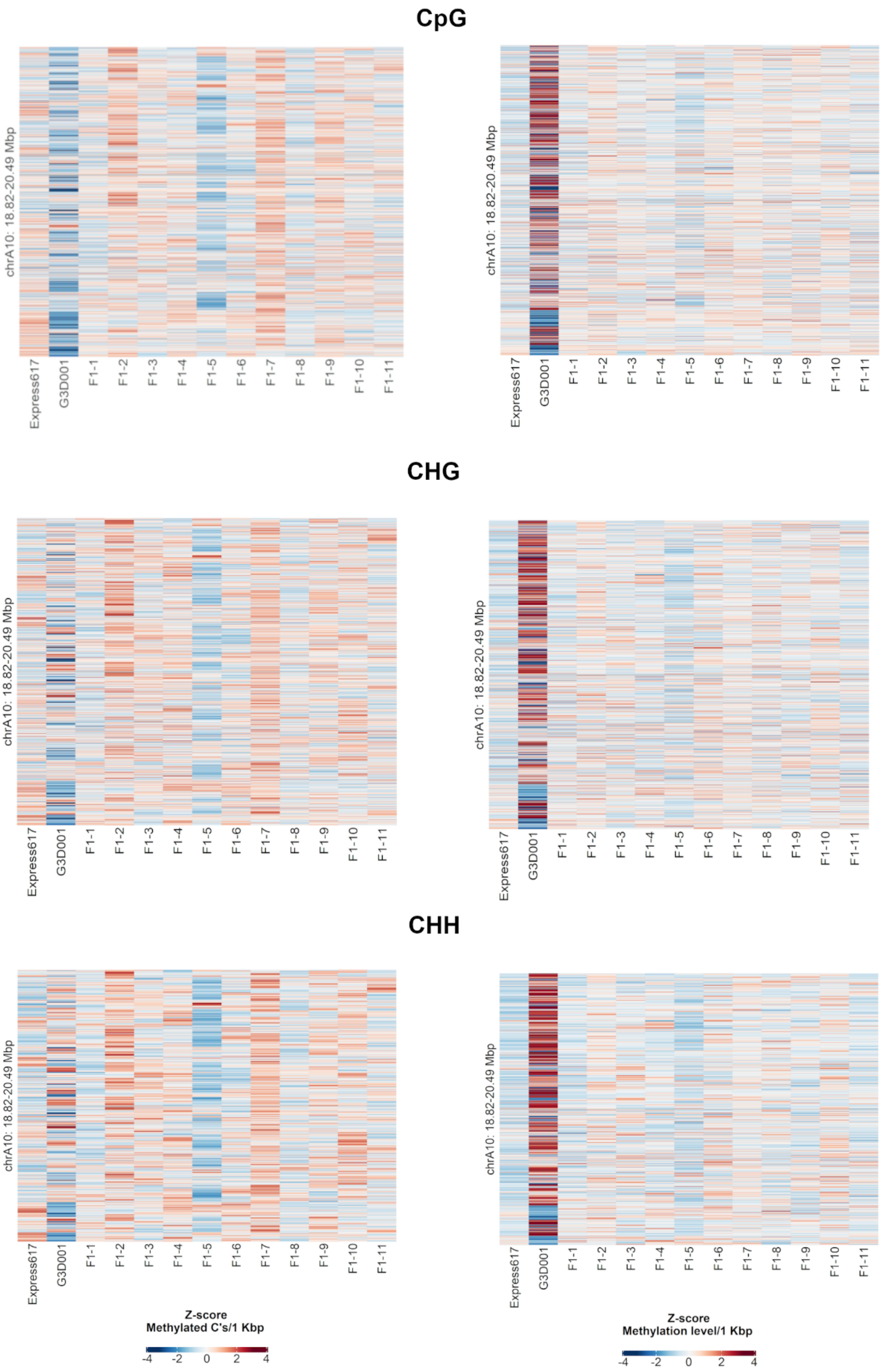


**Supplementary Figure 20. Count of methylated cytosines (left) and methylation level (right) per methylation context within 1 kbp bins across F1 sister plants and parents in chromosome A10: 18.82– 20.49 Mbp.** Bins are sorted from bottom to top from heatmaps by ascending genomic position.

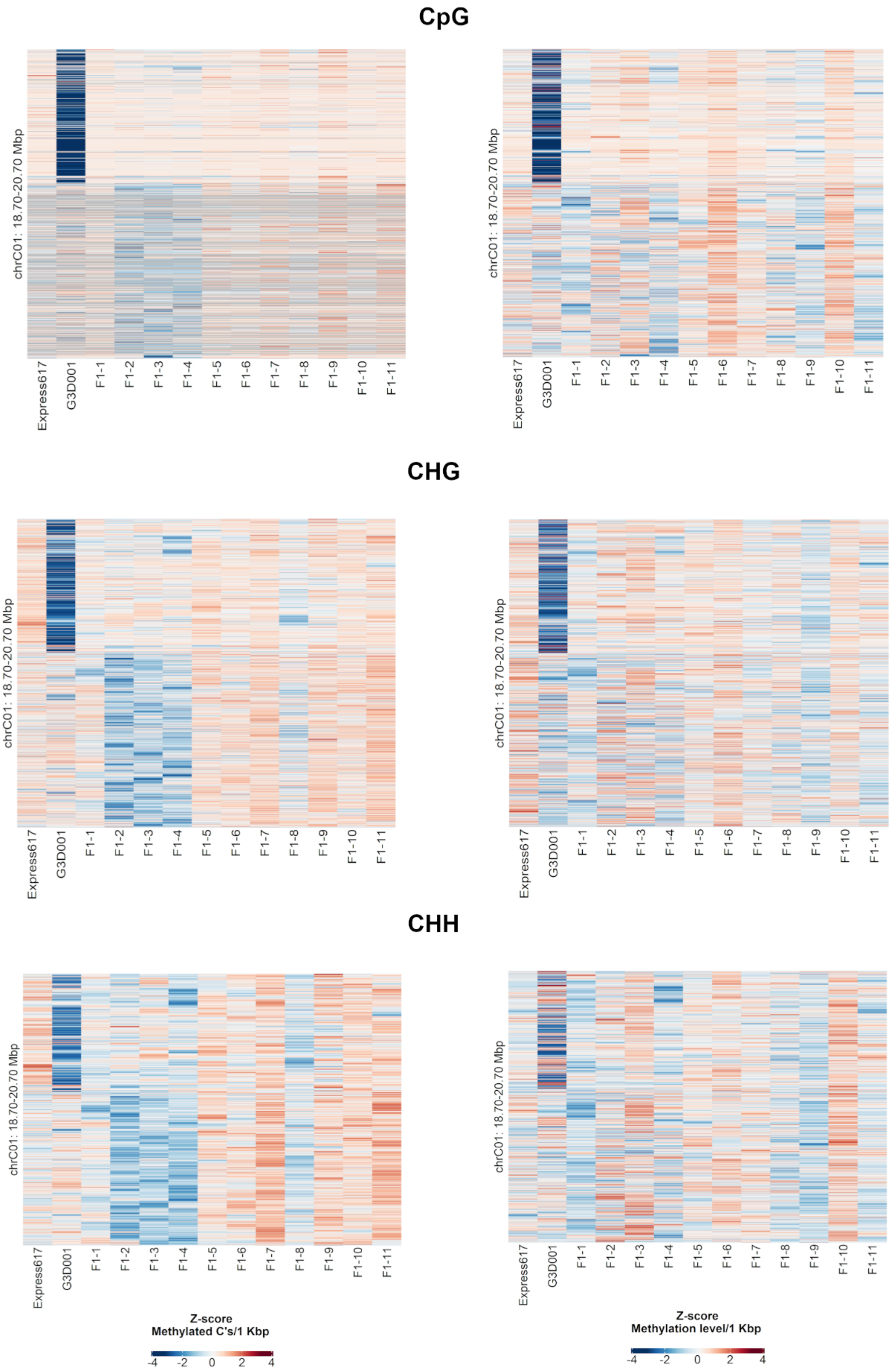


**Supplementary Figure 21. Count of methylated cytosines (left) and methylation level (right) per methylation context within 1 kbp bins across F1 sister plants and parents in chromosome C01: 18.7– 20.7 Mbp.** Bins are sorted from bottom to top from heatmaps by ascending genomic position.

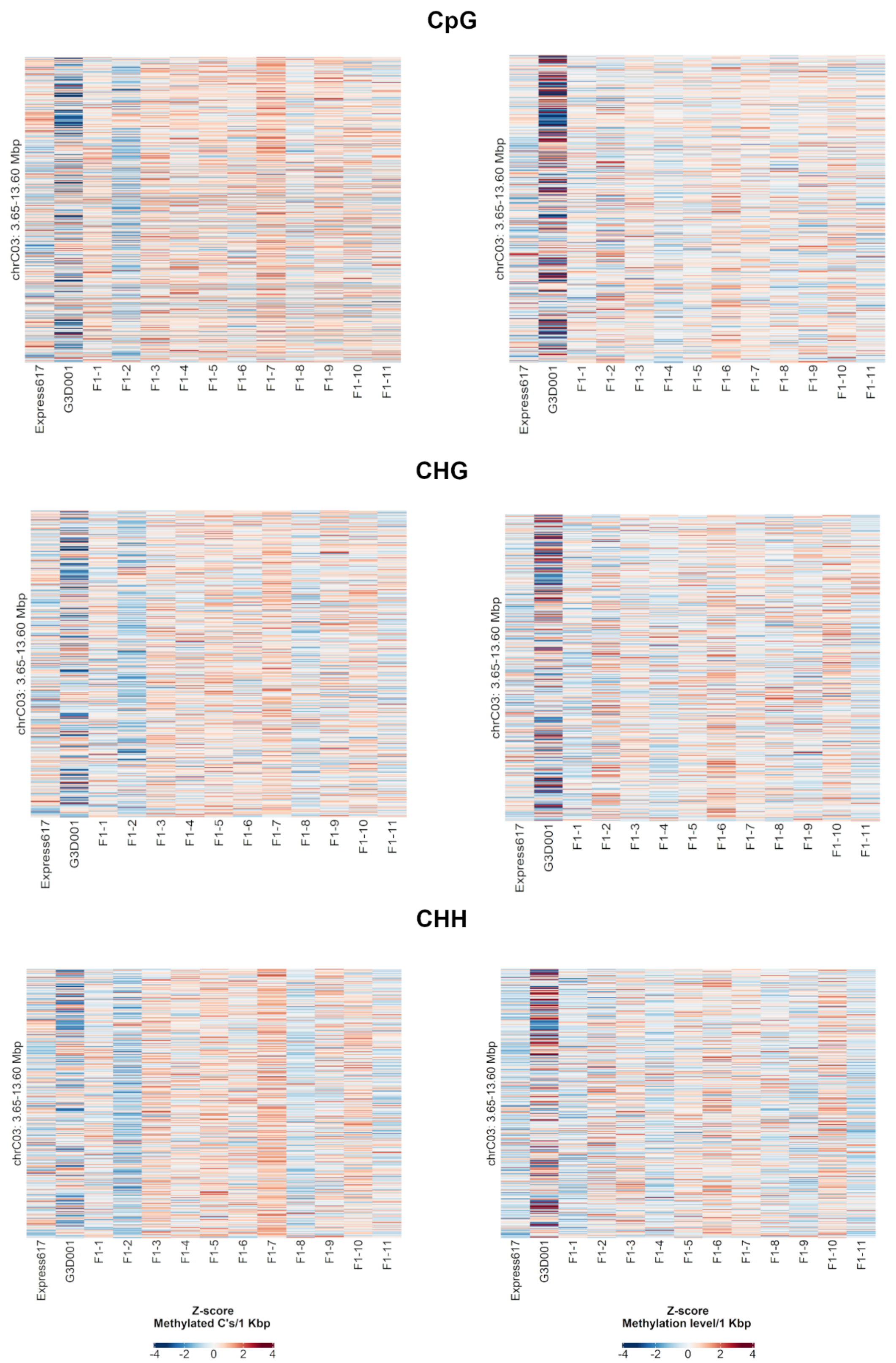


**Supplementary Figure 22. Count of methylated cytosines (left) and methylation level (right) per methylation context within 1 kbp bins across F1 sister plants and parents in chromosome C03: 3.65– 13.60 Mbp.** Bins are sorted from bottom to top from heatmaps by ascending genomic position.

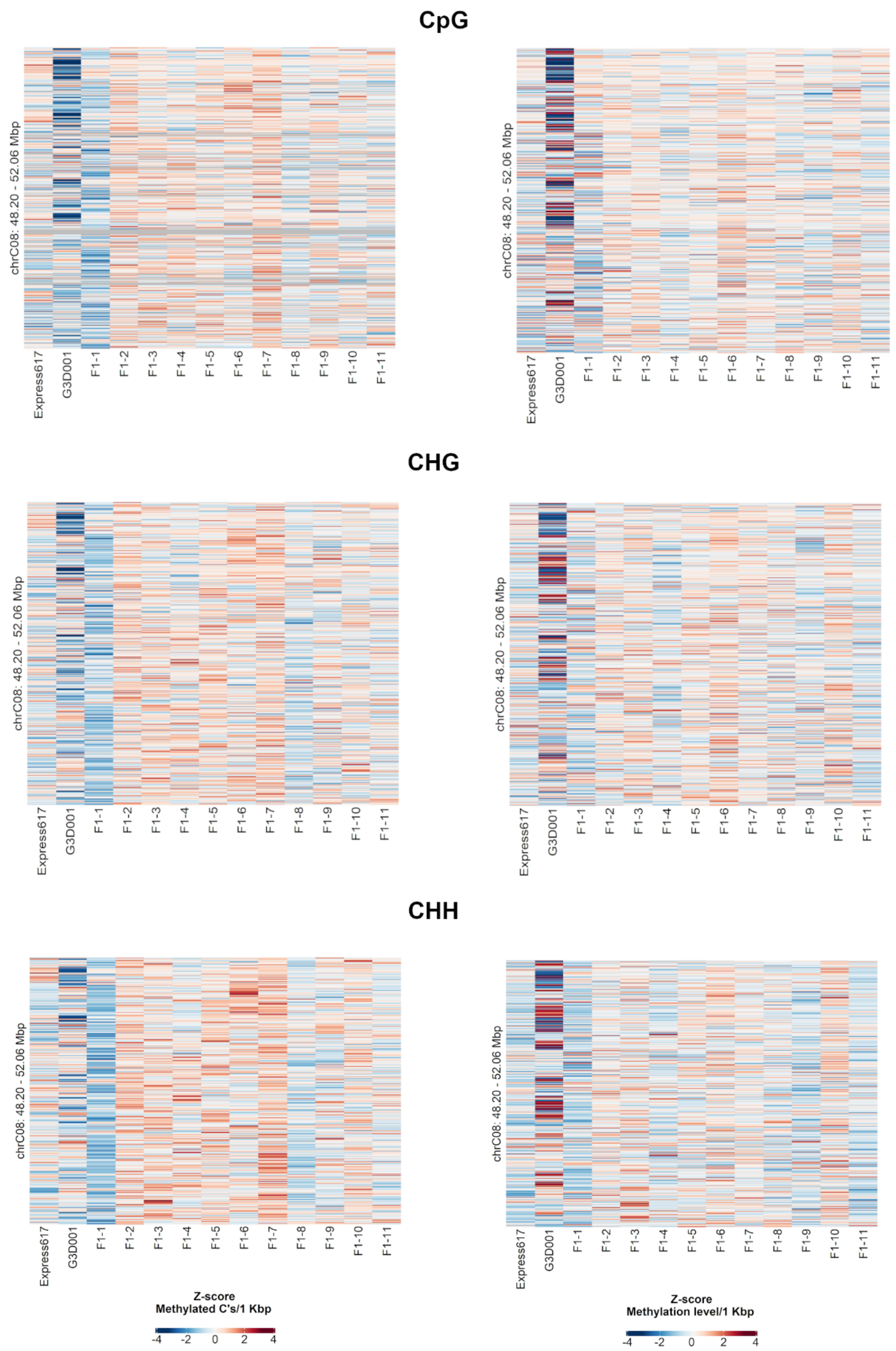


**Supplementary Figure 23. Count of methylated cytosines (left) and methylation level (right) per methylation context within 1 kbp bins across F1 sister plants and parents in chromosome C08: 48.2– 52.06 Mbp.** Bins are sorted from bottom to top from heatmaps by ascending genomic position.

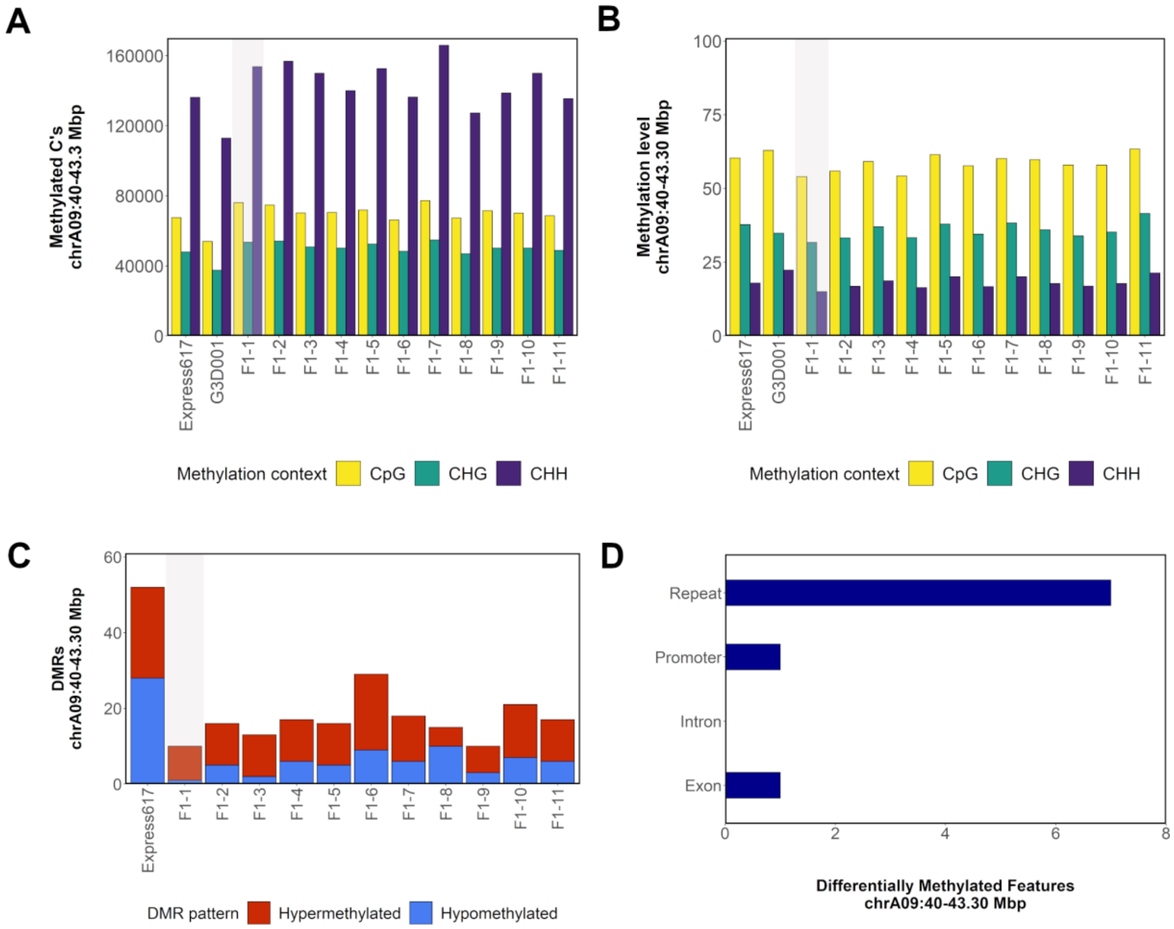


**Supplementary Figure 24. Methylation patterns in chromosome A09: 40– 43.3. a**, Count of methylated cytosines per methylation context. **b**, Methylation level per methylation context. **c**, Count of hypo- and hypermethylated DMRs in comparison to G3D001. **d**, Distribution of DMRs across introns, exons, repeats and promoters (1 kbp upstream from gene start). The number in the F1 samples indicates their biological replicate name. A genotype carrying a spontaneous NRHE on chromosome A09 is highlighted in gray.

.

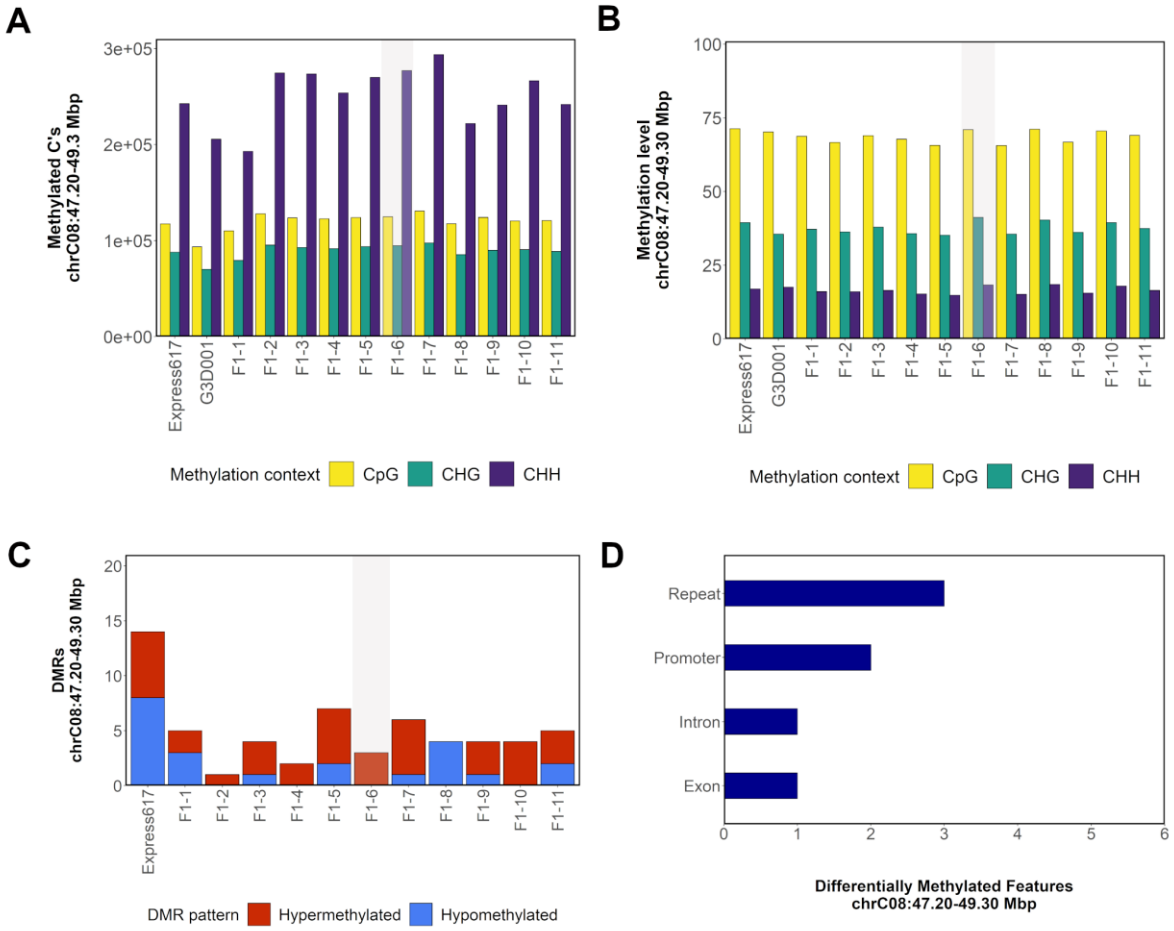


**Supplementary Figure 25. Methylation patterns in chromosome C08: 47.2– 49.3. a**, Count of methylated cytosines per methylation context. **b**, Methylation level per methylation context. **c**, Count of hypo- and hypermethylated DMRs in comparison to G3D001. **d**, Distribution of DMRs across introns, exons, repeats and promoters (1 kbp upstream from gene start). The number in the F1 samples indicates their biological replicate name. A genotype carrying a spontaneous NRHE on chromosome C08 is highlighted in gray.

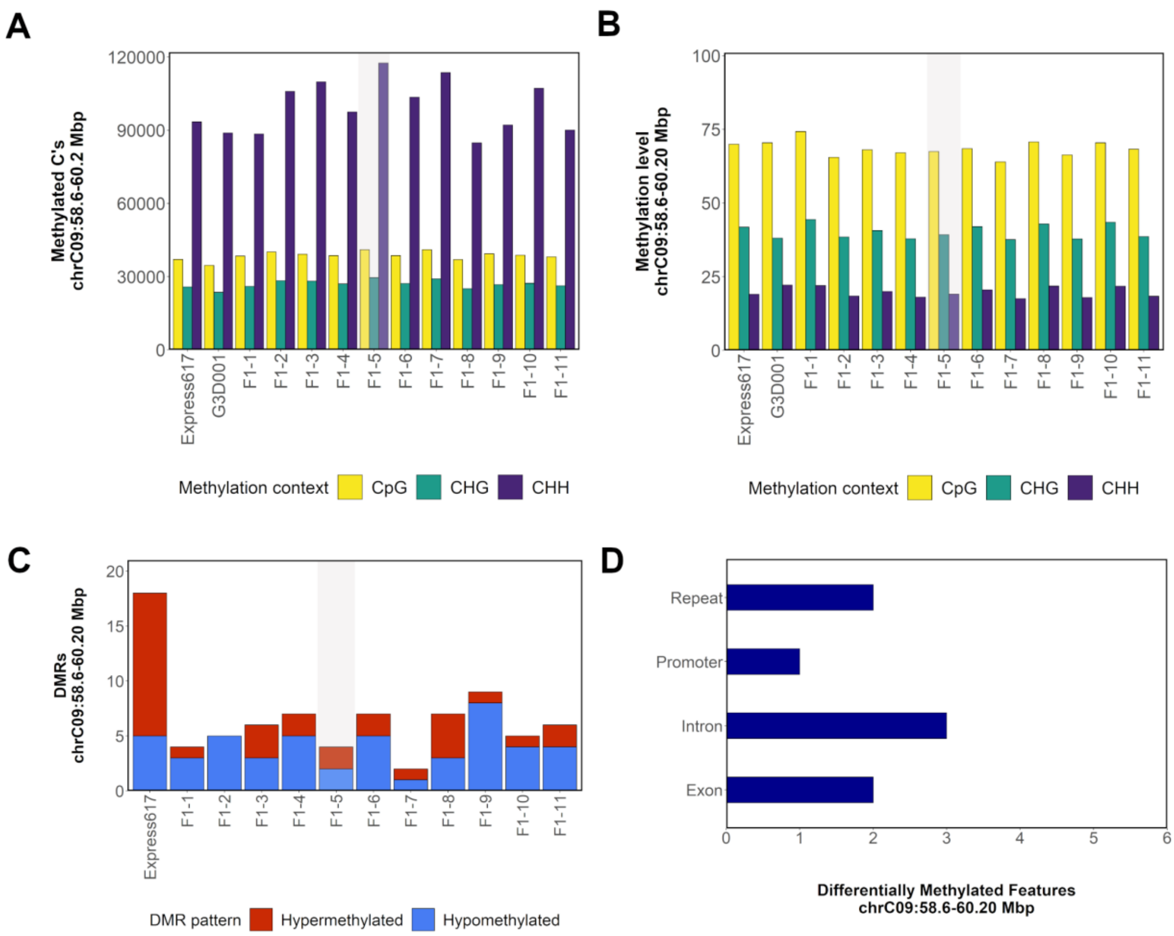


**Supplementary Figure 26. Methylation patterns in chromosome C09: 58.6– 60.2. a**, Count of methylated cytosines per methylation context. **b**, Methylation level per methylation context. **c**, Count of hypo- and hypermethylated DMRs in comparison to G3D001. **d**, Distribution of DMRs across introns, exons, repeats and promoters (1 kbp upstream from gene start). The number in the F1 samples indicates their biological replicate name. A genotype carrying a spontaneous NRHE on chromosome C09 is highlighted in gray.

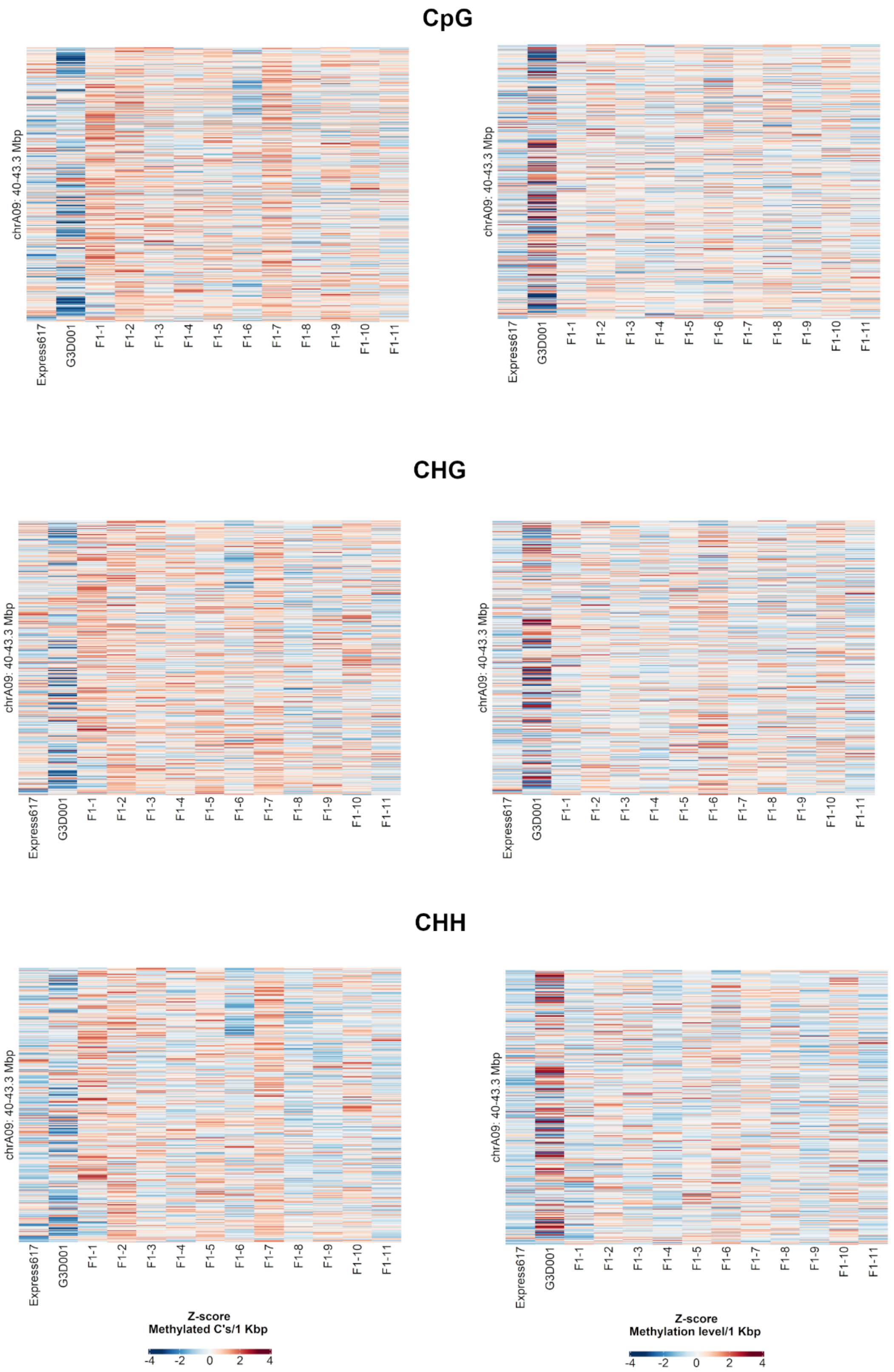


**Supplementary Figure 27. Count of methylated cytosines (left) and methylation level (right) per methylation context within 1 kbp bins across F1 sister plants and parents in chromosome A09: 40– 43.3 Mbp.** Bins are sorted from bottom to top from heatmaps by ascending genomic position.

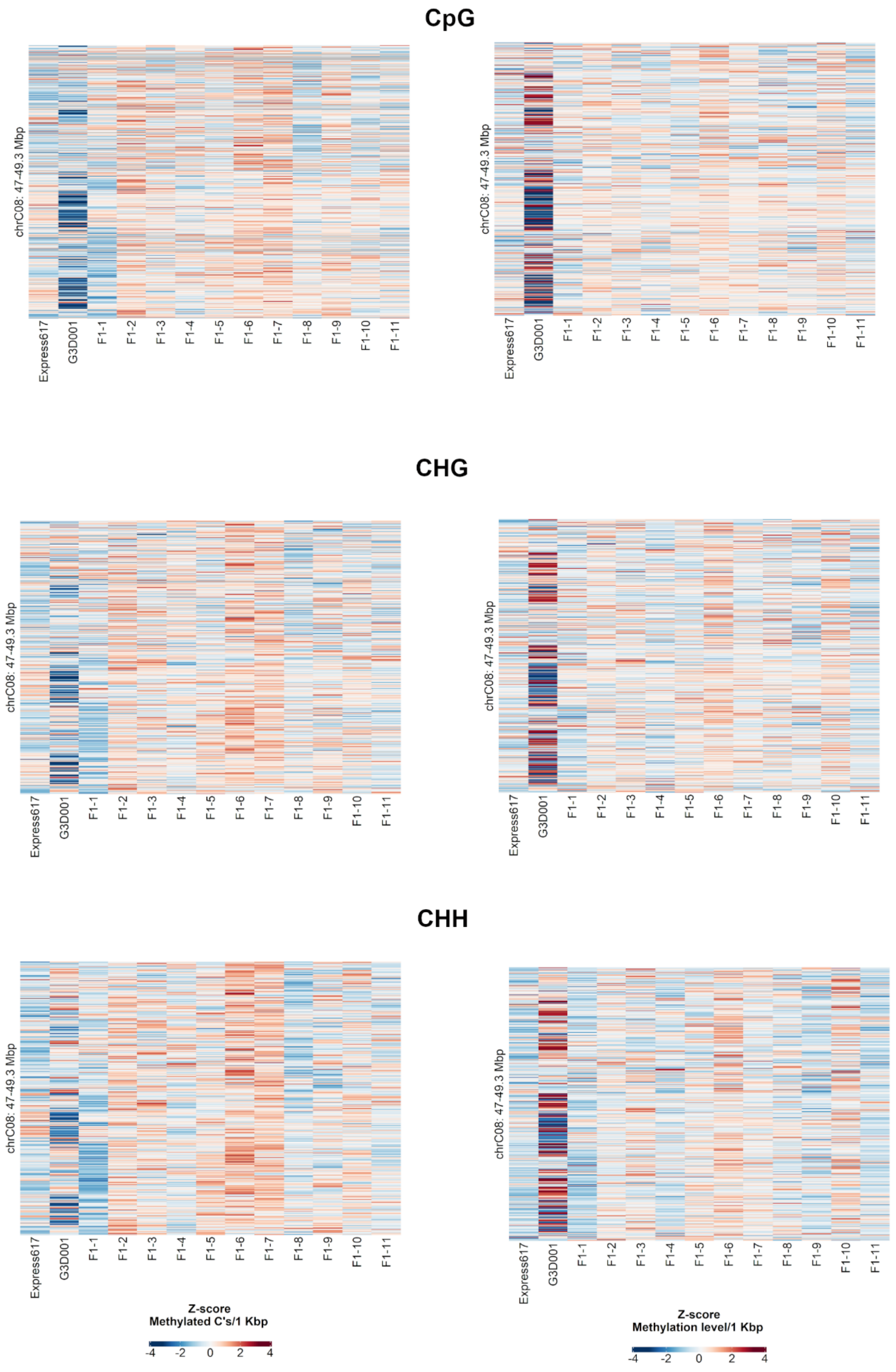


**Supplementary Figure 28. Count of methylated cytosines (left) and methylation level (right) per methylation context within 1 kbp bins across F1 sister plants and parents in chromosome C08: 47.2– 49.3 Mbp.** Bins are sorted from bottom to top from heatmaps by ascending genomic position.

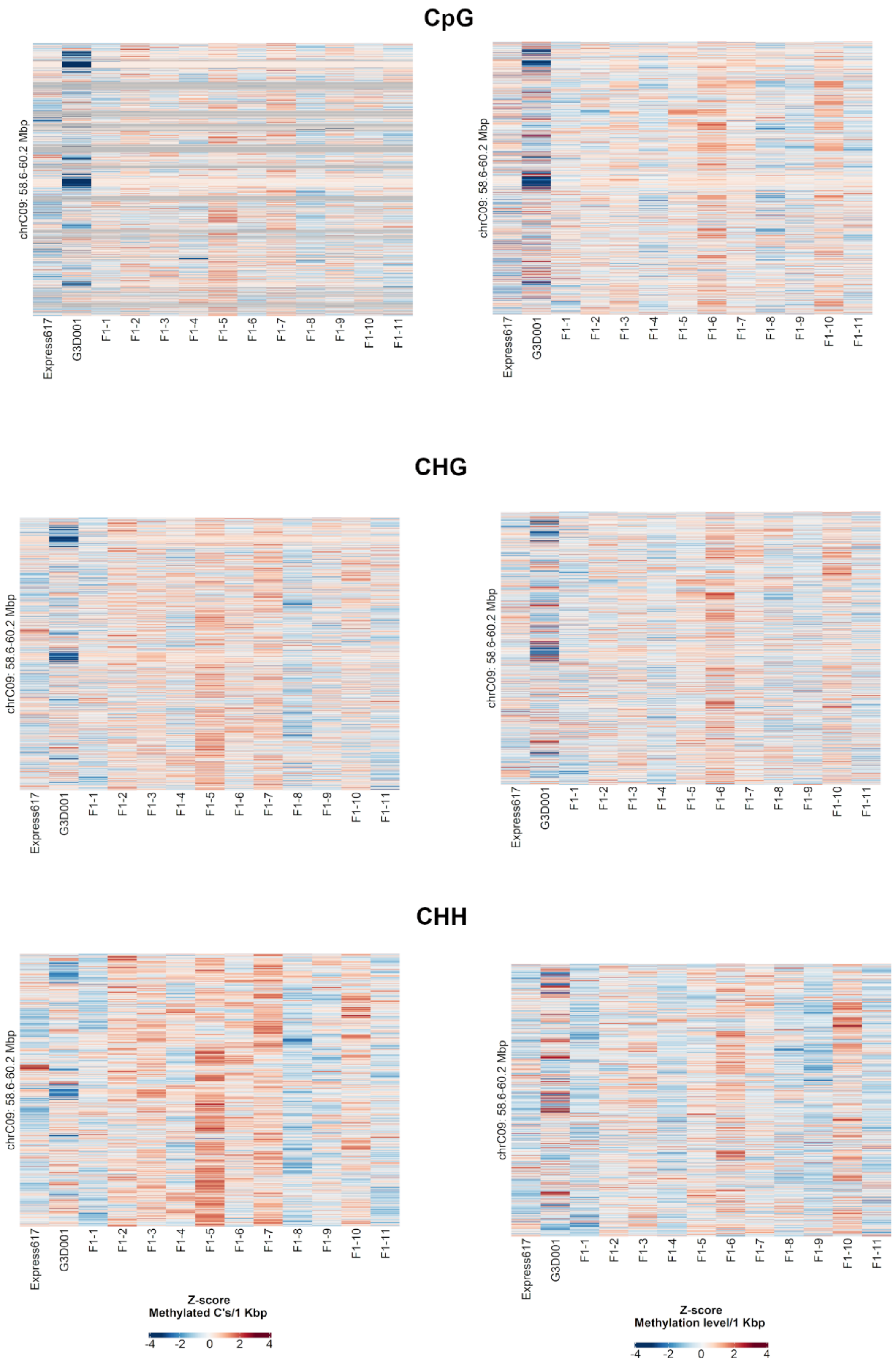


**Supplementary Figure 29. Count of methylated cytosines (left) and methylation level (right) per methylation context within 1 kbp bins across F1 sister plants and parents in chromosome C08: 58.6– 60.2 Mbp.** Bins are sorted from bottom to top from heatmaps by ascending genomic position.
