## Supplementary Data 1 for "Frequent spontaneous structural rearrangements promote rapid genome diversification in a *Brassica napus* F1 generation"

**Supplementary Data 1. Bash script for converting deepsignal split_freq_file_by_5mC_motif python script output to DMRcaller compatible input**

```
#!/bin/bash
for f in `ls fast5s.C.call_mods.frequency_*.CG.tsv | sed 's/fast5s.C.call_mods.frequency_//' | sed 's/.CG.tsv//'`
do
egrep "^chr" fast5s.C.call_mods.frequency_${f}.CG.tsv  | awk -F "\t" '{print $1"\t"$2+1"\t"$3"\t"$7"\t"$8"\tCG"$11}' | sed -e 's/CG../CG\t/g' > prefilt.${f}
done

for f in `ls fast5s.C.call_mods.frequency_*.CHG.tsv | sed 's/fast5s.C.call_mods.frequency_//' | sed 's/.CHG.tsv//'`
do
egrep "^chr" fast5s.C.call_mods.frequency_${f}.CHG.tsv  | awk -F "\t" '{print $1"\t"$2+1"\t"$3"\t"$7"\t"$8"\tCHG"$11}' | sed -e 's/CHG../CHG\t/g' >> prefilt.${f}
done

for f in `ls fast5s.C.call_mods.frequency_*.CHH.tsv | sed 's/fast5s.C.call_mods.frequency_//' | sed 's/.CHH.tsv//'`
do
egrep "^chr" fast5s.C.call_mods.frequency_${f}.CHH.tsv  | awk -F "\t" '{print $1"\t"$2+1"\t"$3"\t"$7"\t"$8"\tCHH"$11}' | sed -e 's/CHH../CHH\t/g' >> prefilt.${f}
done

for f in `ls prefilt.* | sed 's/prefilt.//'`
do
sort -t $'\t' -k 1,1 -k 2,2n prefilt.${f} > filt.${f}.tsv
done
```
